# Kinetics of Xist-induced gene silencing can be predicted from combinations of epigenetic and genomic features

**DOI:** 10.1101/510198

**Authors:** Lisa Barros de Andrade e Sousa, Iris Jonkers, Laurène Syx, Julie Chaumeil, Christel Picard, Benjamin Foret, Chong-Jian Chen, John T. Lis, Edith Heard, Edda G. Schulz, Annalisa Marsico

**Affiliations:** Otto Warburg Laboratories, Max Planck Institute for Molecular Genetics, Ihnestraße 63, 14195 Berlin, Germany; Department of Molecular Biology and Genetics, Cornell University, 416 Biotechnology Building, 14853, Ithaca, New York, USA; Institut Curie, PSL Research University, CNRS UMR3215, INSERM U934, UPMC Paris-Sorbonne, 26 Rue d’Ulm, 75005 Paris, France; Institut Curie, PSL Research University, Mines Paris Tech, INSERM U900, 75005 Paris, France; Department of Mathematics and Informatics, Free University of Berlin, Arnimallee 14, 14195 Berlin, Germany; Annoroad Gene Technology Co., Ltd, Beijing, China; Department of Genetics, University Medical Centre Groningen, University of Groningen, 9700 RB Groningen, The Netherlands; Institut Cochin, Inserm U1016, CNRS UMR8104, Université Paris Descartes-Sorbonne Paris Cité, Paris, France.

**Keywords:** X chromosome inactivation, Xist-mediated gene silencing kinetics, PRO-Seq, Random Forest modelling, silencing pathways

## Abstract

To initiate X-chromosome inactivation (XCI), the long non-coding RNA Xist mediates chromosome-wide gene silencing of one X chromosome in female mammals to equalize gene dosage between the sexes. The efficiency of gene silencing, however is highly variable across genes, with some genes even escaping XCI in somatic cells. A genes susceptibility to Xist-mediated silencing appears to be determined by a complex interplay of epigenetic and genomic features; however, the underlying rules remain poorly understood. We have quantified chromosome-wide gene silencing kinetics at the level of the nascent transcriptome using allele-specific Precision nuclear Run-On sequencing (PRO-seq). We have developed a Random Forest machine learning model that can predict the measured silencing dynamics based on a large set of epigenetic and genomic features and tested its predictive power experimentally. While the genomic distance to the Xist locus is the prime determinant of the speed of gene silencing, we find that also pre-marking of gene promoters with polycomb complexes is associated with fast silencing. Moreover, a series of features associated with active transcription and the O-GlcNAc transferase Ogt are enriched at rapidly silenced genes. Our machine learning approach can thus uncover the complex combinatorial rules underlying gene silencing during X inactivation.

## Introduction

X-chromosome inactivation (XCI) is a developmental process in mammals that ensures equal gene dosage of X-linked genes between XX and XY individuals by transcriptional inactivation of one of the two X chromosomes in female cells (Galupa and Heard 2015). In placental mammals XCI is triggered by the long non-coding (lncRNA) RNA Xist, which is transcribed from a genomic region called the X-inactivation center (*Xic*). *Xist* is upregulated in a monoallelic fashion and its RNA coats the future inactive X chromosome in *cis* leading to the recruitment of several factors involved in transcriptional inactivation and eventually converting the entire X chromosome into silent heterochromatin (Escamilla-Del-Arenal et al. 2011; Gendrel and Heard 2014; Galupa and Heard 2015).

Early events following Xist coating of the chromosome include the depletion of RNA polymerase II from the Xist RNA domain and loss of active histone marks as well as gain of repressive chromatin modifications, such as H2AK119ub1 and H3K27me3, deposited by the polycomb repressive complexes (PRC) 1 and 2 respectively. Later chromatin modifications become associated with the X undergoing XCI, including accumulation of the histone variant macroH2A and DNA methylation of gene promoters (Escamilla-Del-Arenal et al. 2011; Gendrel and Heard 2014; Galupa and Heard 2015). The manner in which Xist RNA induces gene silencing and chromatin changes is still poorly understood, although progress has been made in identifying some of its partners and also the parts of this lncRNA that mediate its different functions. *Xist* contains multiple conserved repeats. The most highly conserved A repeat is thought to mediate gene silencing through recruitment of Spen and other factors including Rbm15 (Chu et al. 2015; McHugh et al. 2015). Repeat-B is required for recruitment of polycomb-repressive complexes mediated by hnRNPK (Pintacuda et al. 2017). While the A-repeat pathway seems to be required for Xist mediated silencing in mouse embryonic stem cells (mESCs) (Wutz et al. 2002), it has been shown to be partially dispensable in extraembryonic tissues, such that a subset of X-linked genes can still be silenced by an Xist mutant lacking the A repeat (Sakata et al. 2017). The PRC1/2 pathway is particularly important for the early maintenance of XCI in extraembryonic tissues but also also seems to contribute to some degree of silencing in epiblast/embryonic stem cells (Wang et al. 2001; Brockdorff 2017).

Intriguingly, the dynamics of Xist-mediated silencing is known to be highly variable between genes across the X chromosome (Chow et al. 2010; Borensztein et al. 2017), with a subset of so-called escape genes remaining active even in somatic cell (Berletch et al. 2011). However, the determinants of susceptibility to XCI remain poorly understood. Since XCI is a multi-step process, local interference with any step, such as coating by *Xist* or access to the silencing machinery of one or several silencing pathways, could delay or prevent silencing of a certain gene or genomic region. Defining some of the underlying features that could explain differential susceptibility to XCI remains an important question, particularly as it is becoming clear that some genes that are not fully silenced are implicated in diseases, such as autoimmune syndromes (Bianchi et al. 2012).

The coating by Xist RNA of the X chromosome would seem the most obvious determinant for X-linked gene silencing dynamics and efficiency. Xist RNA spreading is thought to occur by proximity transfer to sites that are close to the *Xist* locus genomically or in 3D space (“Xist entry sites”) (Engreitz et al. 2013). From there *Xist* has been proposed to move first into gene dense regions and then spread to intergenic domains of the X chromosome (Engreitz et al. 2013; Simon et al. 2013). In differentiated cells Xist binding is observed across the entire X chromosome, but is reduced at escape genes (Engreitz et al. 2013; Simon et al. 2013). Xist RNA is positively correlated with gene density and with PRC2 enrichment and negatively correlated with the density of LINE elements (Engreitz et al. 2013; Simon et al. 2013). Similarly to Xist spreading, kinetics of X-linked gene silencing has also been associated with the genomic distance from the *Xist* locus, such that genes at the distal ends of the X chromosome are silenced late or escape (Marks et al. 2015; Borensztein et al. 2017). In addition, more efficient gene silencing has been linked to the presence of LINE elements (Loda et al. 2017) and full length, active LINEs have been found to be enriched in regions that are otherwise prone to escape (Chow et al. 2010). Genes that are silenced efficiently also tend to be enriched for polycomb complexes (Ring1B, H3K27me3) and depleted for active marks such as H3K4me3 and H3K27ac prior to Xist-induced silencing (Kelsey et al. 2015; Loda et al. 2017). Finally, genes that can actively escape from XCI have been reported to be characterised by CTCF at their transcription start sites compared to partially or fully silenced X-linked genes (Loda et al. 2017). Thus, a variety of genetic and epigenetic features have been implicated in controlling gene-specific silencing efficiency. However, none of these features alone can predict whether and to what extent a gene will be silenced upon XCI, and the associations with measured silencing efficiencies are generally weak. Since no predictive pattern of features has so far been identified, the susceptibility of genes to Xist-mediated silencing is likely to be controlled by a complex combination of different features.

In this study we set out to identify the genetic and chromatin features of X-linked loci, or more precisely the enrichment or depletion of certain features, that predispose genes on the chromosome X to be efficiently silenced or avoid silencing. For this, we measured chromosome-wide silencing dynamics of X-linked genes following induction of *Xist* expression, using allele-specific Precision nuclear Run-On sequencing (PRO-seq) (Kwak et al. 2013). We then trained two Random Forest machine-learning models to predict from 74 genomic and epigenetic features 1) whether a gene is subject to XCI and 2) whether it will be silenced with fast or slow kinetics. Through forest-guided gene clustering we identified feature sets that determine the silencing dynamics of sub-groups of genes, indicating that variable silencing efficiencies might be associated with distinct silencing pathways. For example, we identified a subset of rapidly silenced genes that are pre-marked by PRC1 binding, while genes that escape XCI tend to be very distal to the Xist locus and depleted of CpG islands. We have thus developed a framework to comprehensively assess the contribution of genetic and epigenetic factors to transcriptional silencing on chromosome X in an unbiased and quantitative manner.

## Results

### Quantification of gene-specific silencing dynamics by PRO-seq

To explore the features that control susceptibility to XCI for individual genes, we used an experimental system that accurately measures Xist-induced silencing over time in a chromosome-wide manner. Although differentiating mouse embryonic stem cells (mESCs) are the classical *in vitro* system used to study XCI, this model is not optimal for analysis of gene silencing kinetics as Xist up-regulation and thus gene silencing only occurs in a subset of cells (maximum 60-70%, sometimes as low as 10-20%) and in a rather asynchronous manner, due to cell-to-cell variability in differentiation speed (Chow et al. 2010). This results in a very poor temporal resolution of population measurements. To circumvent this limitation, we used the doxycycline-responsive female TX1072 mESC line (Schulz et al. 2014), where Xist upregulation from the endogenous locus on one X chromosome can be induced by doxycycline treatment in undifferentiated cells in a fast and efficient manner. To assess the dynamics of gene silencing we measured the nascent transcriptome by Precision nuclear Run-on sequencing (PRO-seq) (Kwak et al. 2013) before doxycycline treatment and at 7 different time points after treatment, namely at 0.5, 1, 2, 4, 8, 12 and 24h (Figure 1a). Since the TX1072 line is derived from a cross between two different mouse strains (C57BL/6 x CAST/EiJ), a large number of polymorphisms enables allele-specific mapping of the PRO-seq reads to the two parental genomes to obtain the number of reads originating from the B6 or Cast X-chromosomes, respectively. The data was highly reproducible, since replicates generated for the first and last time point of the experiment (0h, 24h) were strongly correlated (pearson correlation > 0.94; **Supplemental Figure S1**). *Xist* started to be upregulated from the B6 chromosome about 1h after Dox treatment and reached a plateau after 4h (Figure 1b, Figure 1c and **Supplemental Figure S2**). Global expression of the B6 X chromosome, which carries the doxycycline-inducible promoter was gradually reduced over time due to X inactivation, starting at 4 hours of treatment (Figure 1d). Based on the PRO-seq dataset we estimated gene-specific silencing half-times in a chromosome-wide manner, covering 296 genes in total.

**Figure 1:**
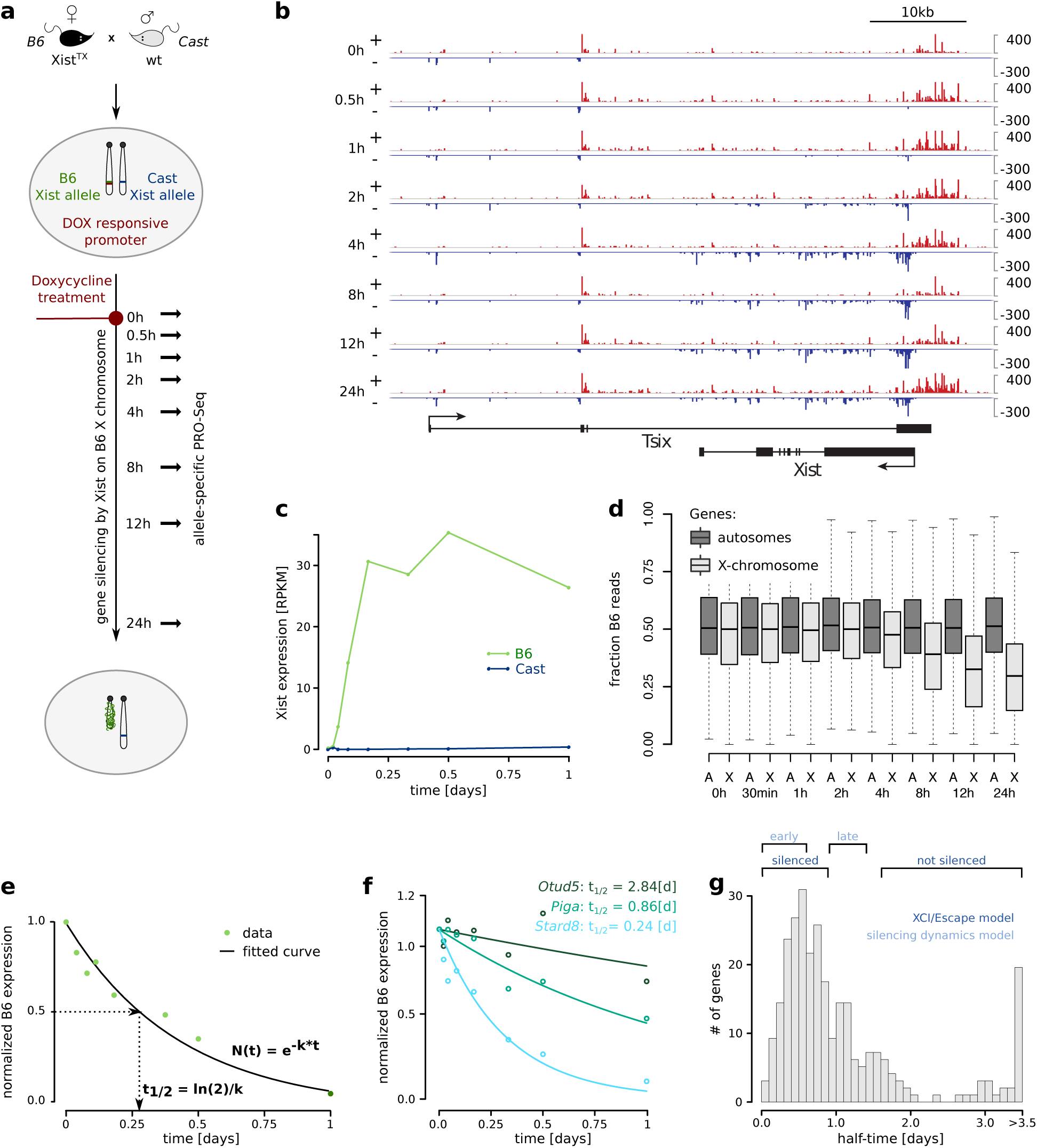
Measuring gene silencing dynamics. (**a**) Schematic overview of the experimental setup. Generation of hybrid female mESC line (B6:Cast) with non-random XCI due to the insertion of a DOX-responsive promoter in front of the Xist gene on the B6 allele. Pol II activity at base-pair resolution was measured by allele-specific PRO-seq (Precision nuclear Run-On and sequencing assay) in a 24 hours time course. (**b**) Strand-specific read density at the Tsix/Xist locus is shown over time after Dox-mediated induction of Xist. Plus-strand is shown in red, minus strand is in blue; the y-axis is in reads per million. (**c**) Xist kinetics on B6 and Cast over 24 hours time course. (**d**) Distribution of the fraction of B6 reads from the PRO-seq experiment for both autosomal (A) and X-linked (X) genes over time. (**e**) Schematic overview of the estimation of gene silencing half-times from the allele-specific PRO-seq time course data. Raw reads are normalized to the uninduced control and corrected for basal skewing towards one allele. Normalized read counts are fitted to an exponential decay function separately for each gene and the silencing half-time of the corresponding gene is computed. (**f**) Examples of three fitted exponential functions, and corresponding computed half-times, for three genes: one early silenced (*Stard8*), one intermediated silenced (*Piga*) and one known escapee gene (*Otud5*). (**g**) Distribution of estimated half-times for 296 X-linked genes. The half-time ranges used to define the model classes, silenced and not silenced for the XCI/escape model and early vs late silenced for the silencing dynamics model are also highlighted.

The expectation is that, upon initiation of silencing, X-linked transcript expression will continue to exponentially decrease at constant rate and can be described by an exponential decay function typically used to model transcription inhibition time courses, mRNA decay and mRNA half-lives (Rabani et al. 2011; Lugowski et al. 2018).

To this end, expression from the B6 chromosome for each gene was normalized to the uninduced control to correct for basal skewing towards one allele and was then fitted with an exponential decay model to compute the silencing half-time (see Methods section). The half-time indicates the time point at which transcription of the B6 chromosome is reduced by 50% compared to the uninduced control (Figure 1e). The estimated half-times ranged from several hours up to several days (Figure 1f and Figure 1g), showing that not all genes get silenced in the 24h time course that was measured.

### The estimated silencing dynamics are comparable to those measured *in vitro* and *in vivo*

In our experimental design we induced Xist ectopically in undifferentiated mESCs in order to measure gene silencing with high temporal resolution. To ensure that the relative silencing dynamics across genes in this setting are comparable to those in the cellular context where XCI occurs endogenously, we compared the estimated half-times to those measured during differentiation (Xist induction and differentiation for 48h). To this end, we generated two additional data sets again using the TX1072 cell line, where mRNA-Seq was performed at different time points of doxycycline treatment, namely 0, 2, 4, 8, 12 and 24h in undifferentiated mESCs and 0, 8, 16, 14 and 48h in differentiating mESCs (Figure 2a). The computed half-times were very similar whether measured in undifferentiated cells or during differentiation (Figure 2b, Pearson correlation coefficient 0.75, p-value=7.07e-60), suggesting that the differentiation process only has a minor impact on relative gene silencing dynamics. When comparing half-times estimated from the two different data types (mRNA-Seq vs PRO-seq) correlation was generally a bit lower, independent of the cellular context (Figure 2c and Figure 2d, Pearson correlation coefficient 0.52/0.51), which would be expected given that PRO-seq measures the direct transcription dynamics, whereas mRNA-Seq kinetics are modulated by transcription, RNA-processing and degradation. All three datasets measure doxycycline-induced silencing dynamics to ensure inactivation of the same chromosome (B6) in all cells, which is a prerequisite to assess XCI in population measurements. To ensure that doxycycline-induced XCI occurs with similar dynamics as X inactivation in differentiating XX ESCs, we compared our data to a previous study that had used a different, doxycycline-independent strategy to make XCI non-random (Marks et al. 2015). In that study a stop-cassette had been inserted in Xists repressive antisense transcript Tsix, which results in preferential inactivation of the mutant chromosome during differentiation. The silencing classes that had been defined in that study (early, intermediate, late, escapee) are in good agreement with the half-times we have estimated from the PRO-seq data (Figure 2e), suggesting that doxycycline-induced XCI as used in our study recapitulates endogenous gene silencing dynamics.

**Figure 2:**
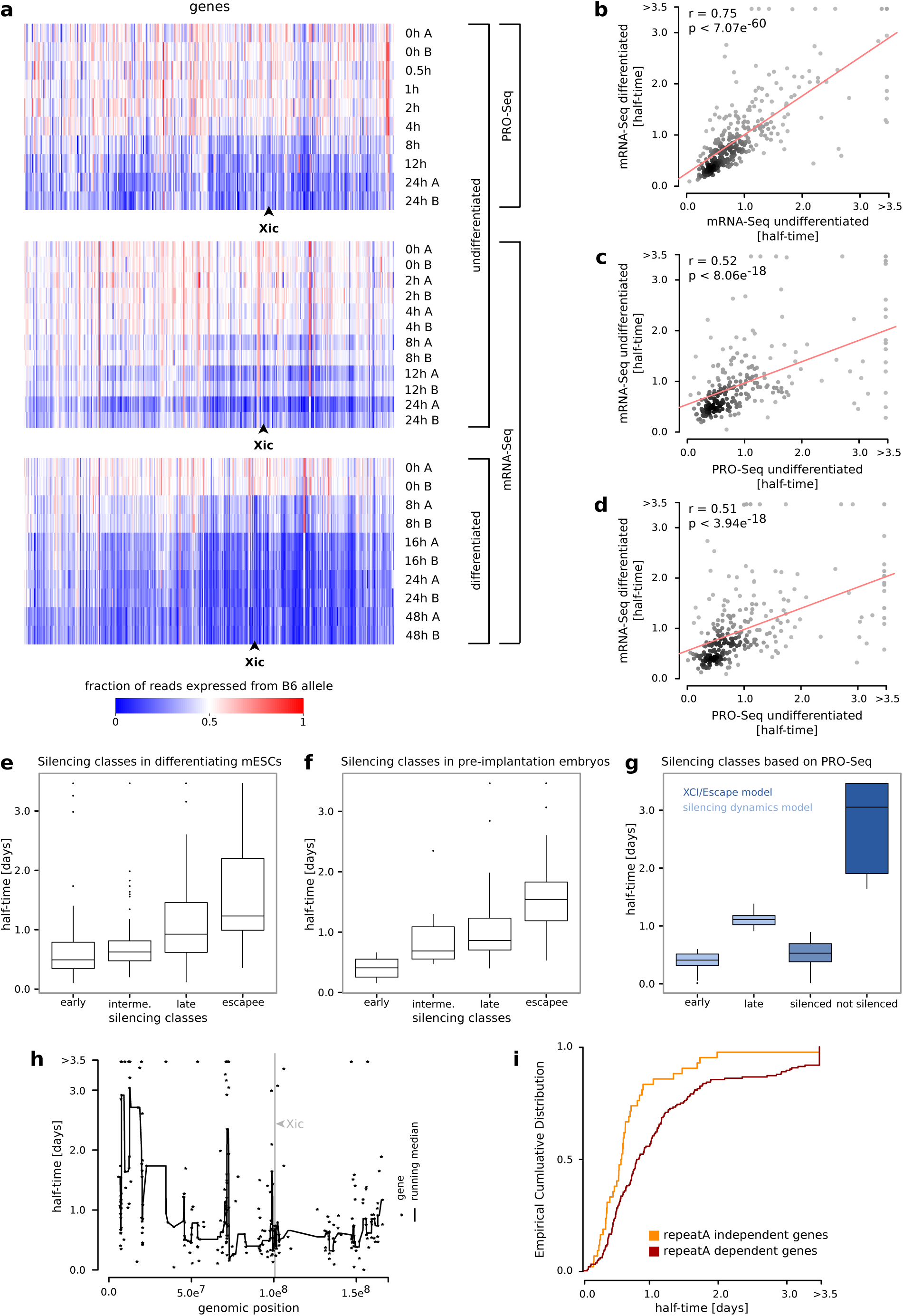
Estimating gene silencing half-times. (**a**) Fraction of reads expressed from B6 for each timestep of the time course experiment ordered by genomic position. X inactivation center is marked with Xic. Upper panel: PRO-seq experiment on undifferentiated mESCs over a time course of 24 hours with replicates for time point 0 and 24 hours; intermediate panel: mRNA-Seq experiment on undifferentiated mESCs over a time course of 24 hours with replicates for each time point; lower panel: mRNA-Seq experiment on differentiated mESCs over a time course of 48 hours with replicates for each time point. (**b - d**) Comparison of half-times computed from PRO-seq data on undifferentiated mESCs, mRNA-Seq data on undifferentiated mESCs and mRNA-Seq data on differentiating mESCs. For each comparison a scatterplot with a fitted regression line (red) is shown. Pearson correlation between half-times and significance of correlation (p-value of test statistics) is reported for each plot. (**e**) Distribution of our computed half-times within the silencing classes defined by Marks et. al. from RNA-seq time course data on differentiating mESCs (Marks et al. 2015). (**f**) Distribution of our computed half-times within the silencing classes defined by Borensztein et. al. during XCI in pre-implantation mouse embryos from single-cell RNA-seq data (Borensztein et al. 2017). (**g**) Distribution of our computed half-times within the classes defined by our classification models: silenced and not silenced (XCI/escape model) and early and late silenced (silencing dynamics model), respectively. (**h**) Estimated half-times (black stars) relative to genomic positions of the corresponding genes. A fitted smooth curve of the half-times is displayed as black line. The X inactivation center is marked with a grey line. **i** Comparison of our estimated half-times with the X-linked gene dynamics observed in an Xist A-repeat mutant cell line (Sakata et al. 2017). “repeatA independent genes” refers to the half-time cumulative distribution of those genes which still undergo silencing in the mutant cell line (orange line), while “repeatA dependent genes” refers to the half-time cumulative distribution of those genes were silencing is abrogated in the mutant cell line.

We also validated the silencing half-times estimated *in vitro* by looking at an *in vivo* situation. We compared our Xist-induced silencing half-times in ESCs to the dynamics of (imprinted) XCI in pre-implantation mouse embryos, measured through single-cell RNA-Sequencing (Borensztein et al. 2017). In that study, genes were classified as silenced early (before the 16-cell stage), intermediate (before the 32-cell stage), late (at the blastocyst stage) or not at all (escapee). This was once more in very good agreement with the silencing half-times estimated from the PRO-seq data (Figure 2f), since *in vivo* early silenced genes have half-times of less than one day, late silenced genes have higher half-times between 0.8-1.5 days, while escapees lie in the upper range of computed half-times of above 1 day. Taken together, these results show that the silencing dynamics computed in our study recapitulate well the dynamics of endogenous X-inactivation both *in vitro* and *in vivo*. Thus, based on our PRO-seq data we defined classes of gene expression dynamics: early or late silenced; and silenced or not silenced (including both escapees and very late silenced genes), as described in the next paragraph.

### Identifying determinants of gene silencing dynamics with Random-Forest modelling

Having estimated gene-specific silencing kinetics from our PRO-seq data we set out to understand which features might determine whether a gene is subject to XCI at all and whether it is silenced with fast or slow kinetics. First, we noted that genes close to the *Xic*, where the *Xist* gene is located, tended to be silenced faster than distal genes (Figure 2h), in agreement with a previous study (Marks et al. 2015). However, many genes did not follow this trend, as they are close to the *Xic* but escape XCI or are located in the distal regions of the X chromosome but are silenced fast. To uncover additional factors that potentially determine the susceptibility to *Xist*-mediated silencing of a given gene we developed a statistical machine learning model to predict silencing dynamics based on genomic and epigenetic features.

We collected more than 100 publicly available high-throughput datasets (ChIP-seq and Bisulfite-Seq) measuring chromatin modifications, chromatin modifiers, transcription factor binding (TF) and components of the transcriptional machinery (Table 1). As these data sets had been generated in undifferentiated mESCs, they correspond to the chromatin state before *Xist* induction. After stringent filtering on data quality (see Methods section, **Supplemental Figure S14** and **S15** for examples), we computed the enrichment for 57 of these ChIP-Seq features at the promoters of all genes present in our PRO-seq data set (**Supplemental Table S5**). The active transcription start site (TSS) of each gene was identified based on a bidirectional PRO-seq pattern (Core et al. 2014; Danko et al. 2015) (**Supplemental Figure S3** and Methods section). Out of 296 gene for which we computed half-times 280 could be assigned an active TSS and were used for subsequent analysis. In addition, we included a series of genomic and structural features, such as gene density, the frequency of 3D chromatin interactions with different genomic elements and the linear distance to other genomic features, such as the *Xist* locus, the next TAD boundary and the next lamin-associated domain (LAD) (Table 1, for further details on the collected data and for data pre-processing refer to Methods section and **Supplemental Text S1**).

**Table 1:** Epigenetic and genomic features used for modeling.

|  |  |  |
| --- | --- | --- |
| epigenetic features | sequence-specific transcription factors | cMyc, Esrrb, Klf4, MafK, Nanog, nMyc, Oct4, Sox2, Tcf3, Tcfcp2l1, YY1, Znf384 |
|  | structural proteins | CTCF, Smc1, Smc3 |
|  | general transcription regulators | Cdk9, E2F1, Hcfc1, Max, Med1, Med12, Nipbl, RNAPII (pS2, pS5, pS7, unphosphorylated), Sin3a, Taf1, Taf3, Tbp |
|  | chromatin modifications (activation) | H3K27ac, H3K9ac, H3K4me1, H3K4me3, H3K36me3, H3K79me2, KMT2b/Mll2 |
|  | chromatin modifications (repression) | H2AK119ub1, H3K27me3, Ring1b (PRC1), Cbx7 (PRC1) Rybp (PRC1), KMT6/Ezh2 (PRC2), Suz12 (PRC2), Kdm1A/Lsd1, Kdm2A, Kdm2B, Hdac1, Hdac2, DNA methylation (BS-Seq), 5fC (MeDip), 5hmC (MeDip), Tet1 |
|  | others | H2A.Z, Ogt, Brg1, Cbx3 |
| genomic features | genomic elements | distance to the Xist locus<br>overlap with Xist entry sites<br>distance to TAD border<br>distance to LAD |
|  |  | overlap with LAD<br>gene density (200kb)<br>overlap with CpG islands<br>CpG content |
|  | 3D structure | number interactions (HiC) all,<br>mean interaction strength (HiC) all, number interactions (HiC) promoter,<br>mean interaction strength (HiC) promoter, mean interaction strength (HiC) xist, number interactions (HiCap) promoter, number interactions (HiCap) enhancer, number interactions (HiCap) all |

To identify features that determine a genes susceptibility to Xist-mediated silencing, we first attempted to develop a linear model that would predict the silencing half-time from the collected epigenetic and genomic features (data not shown). This simple approach however, had little predictive power, probably because no single linear combination of features or rules could be defined to discriminate, for example, silenced from not silenced genes. The different functional domains of Xist might recruit distinct silencing complexes (e.g. PRC1 and Spen/Hdac3) and thus elicit several parallel silencing pathways. Susceptibility to each pathway might be determined by distinct feature patterns, thus resulting in different sets of rules underlying Xist-mediated XCI. We expected to identify feature combinations associated with different silencing pathways from our data set. For example in the trophoblast, genes that require the Xist A-repeat, which recruits Spen, exhibit longer silencing half-times compared to A-repeat independent genes that might be preferentially targeted by the polycomb-repressive complexes (Sakata et al. 2017) (Figure 2i, Kolmogorov Smirnov (KS) test, p = 2.2*10-6).

To identify different combinatorial rule sets that could predict silencing susceptibility, we chose to use Random Forest (RF), a non-parametric machine learning method which has been shown to perform very well in non-linear classification tasks (Polak et al. 2015). RFs combine an ensemble of single classification trees, which successively split the feature input space in a non-linear fashion, to predict the value of a discrete binary variable (Figure 3).

**Figure 3:**
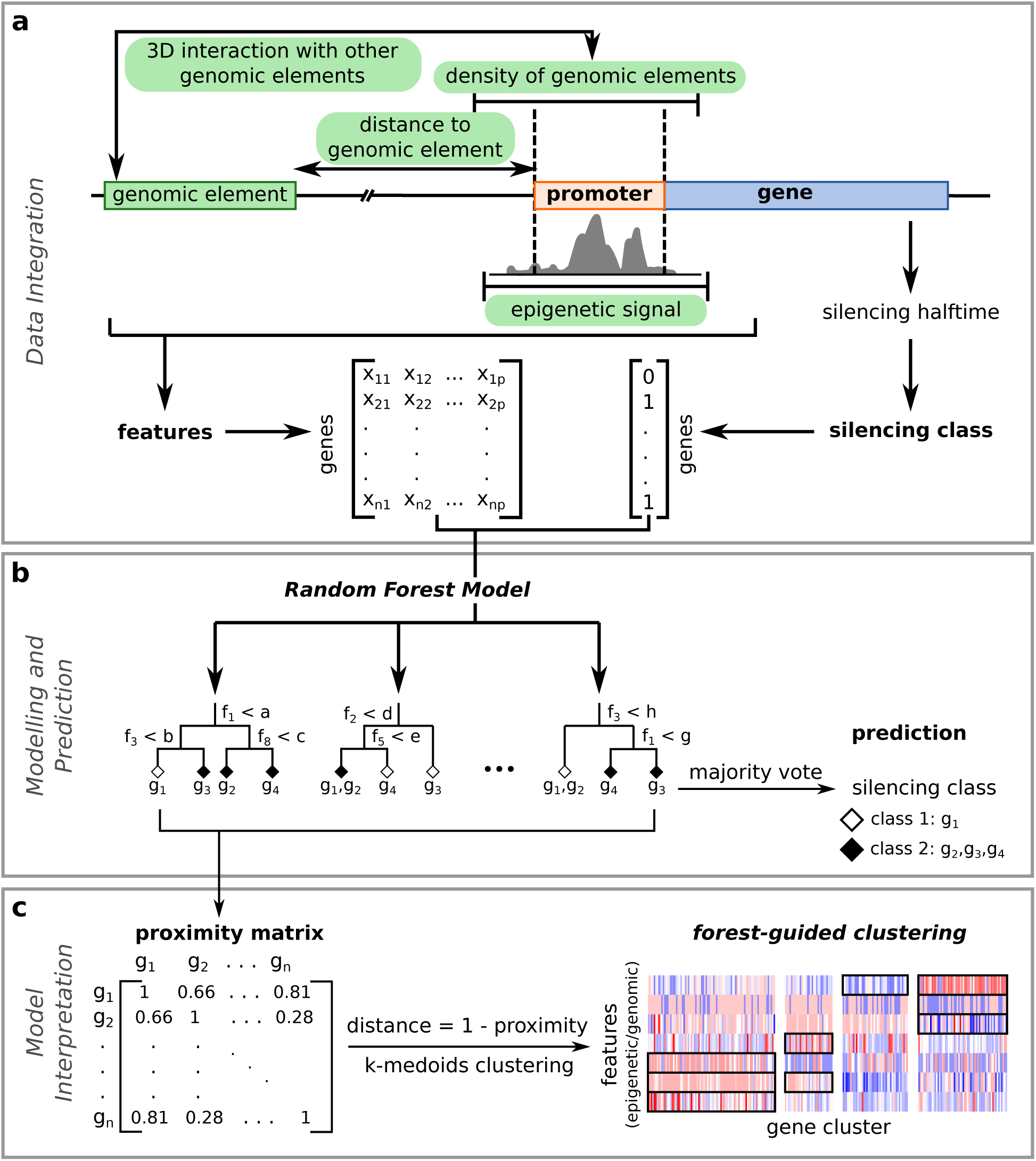
Schematic overview of our modeling approach. (**a**) Input data for the model are collected and pre-processed. In particular, signals from epigenetic modifiers, histone marks and other factors at X-linked gene promoters are computed, as well as features such as distance of genes to other genomic elements (e.g. Xist locus, LADs), density of genomic elements around gene promoters or 3D interactions. Estimated gene half-times are converted to discrete classes for modeling. (**b**) Classification model. Genes are classified either into silenced versus not silenced (XCI/escape model) or early versus late silenced (silencing dynamics model) using in both cases a Random Forest classifier. (**c**) Forest-guided clustering for model interpretation. A similarity matrix between genes (also referred to as proximity matrix) is computed from the trained model and converted into a distance matrix for clustering. Genes that end up more often in the same leaves of the Random forest trees are more likely to be clustered together according to the common subset of features that determine their classification. Clustering results, displaying genes and their most significant associated features are displayed as a heatmap.

Based on the PRO-seq derived silencing half-times, we classified all genes according to whether they are subject to XCI or escape (Figure 1g, silenced/not silenced) and whether they are silenced with slow or fast kinetics (Figure 1g, early/late). The optimal half-time cutoffs were determined during model training (see Methods section and **Supplemental Table S6**). As described above, the resulting classes largely agree with those previously defined in differentiating mESCs and in pre-implantation embryos (Marks et al. 2015; Borensztein et al. 2017) (Figure 2e and Figure 2g). Moreover, the “not silenced” class is enriched for known escapees (odd ratio 2.6, p-value 0.0032, Fisher exact test). The discretization of the gene half-times into three non-disjoint classes and the use of two classification models rather than one continuous model are justified by the fact that the computed half-times tend to be noisy and give rather an indication of the silencing trend for each gene than an exact kinetic measure. We built two binary classification models, one for silenced versus not silenced genes (XCI/escape model), and another one for early versus late silenced genes (silencing dynamics model) to predict a genes silencing susceptibility from a total of 74 epigenetic and genomic features. The XCI/escape model would allow us to pinpoint those factors or combinations of factors which are important for silencing in general, while the silencing dynamics model would identify factors that influence the kinetics of gene silencing.

Both RF models are able to predict gene silencing dynamics with error rates of 27% and 30%, which means that 73% and 70% of genes are classified correctly, which is considerably more than a random predictor (computed as described in Methods section). We assessed the individual contribution of genomic and epigenetic features to the classification accuracy via Random Forest variable importance analysis for the positive (silenced / early) and the negative class (not silenced / late) of each model by the Mean Decrease in Accuracy (MDA) (Figure 4). The higher this value, the more important the contribution of the feature to the classification model (see Methods section). We then trained our models on a set of only 10/11 top features (see Methods section), which greatly improved the prediction error rate to 23% (XCI/escape model) and 22% (silencing dynamics model) as shown in **Supplemental Figure S4**. Discarding low-impact noisy features thus appears to greatly improve modeling of X-chromosomal gene dynamics in both settings. Reassuringly, the XCI/escape model correctly predicted several known escapees, such as *Ddx3x, Taf1, Eif2s3x, Pdbc1, Kdm6a, Usp9x* and *Utp14a*, identified in more than one study (Yang et al. 2010; Berletch et al. 2011; Splinter et al. 2011; Calabrese et al. 2012; Wu et al. 2014; Marks et al. 2015).

**Figure 4:**
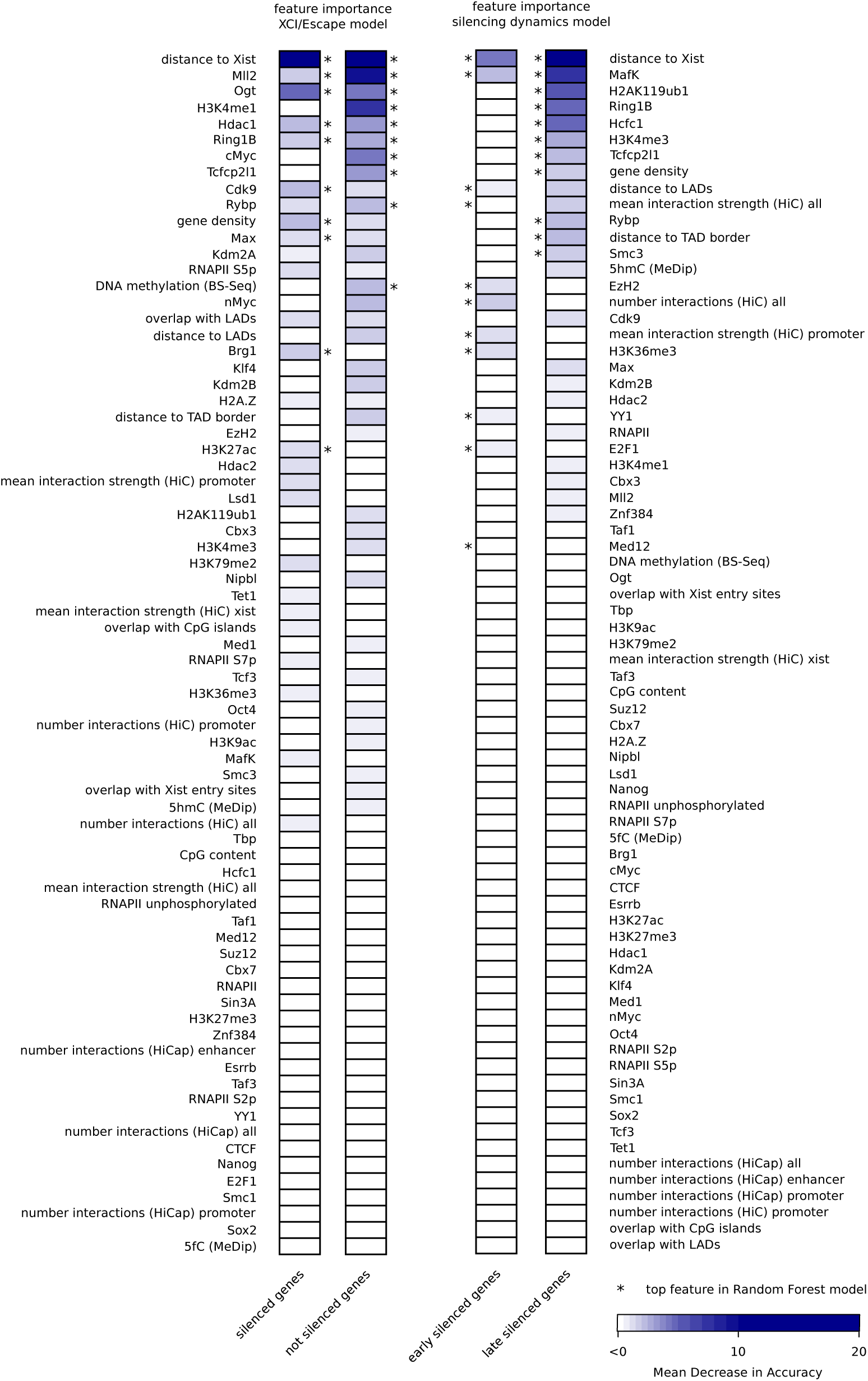
Feature importance for both XCI/escape and silencing dynamics model. For each model features are ranked according to their importance for the classification (either for one class or the other) quantified by the Mean Decrease in Accuracy (MDA) (see Methods section). The top features of each class (10 for XCI/Escape model; 11 for silencing dynamics model) which are used to build the final model are marked with a star. The color gradient follows the MDA values: important features have a high MDA and are shown in dark blue, while completely uninformative features have a low MDA (close to zero or negative) and are shown in white. Intermediate features are shown with different grades of blue. For more details on the stability analysis of important features refer to **Supplemental Figure S5** and **S6**, **Supplemental Text S2**.

The most important feature associated with silencing in both models was close genomic proximity to the *Xist* locus, which has an MDA of 16-21% in the XCI/escape model and of 5-10% in the silencing dynamic model (Figure 4, **Supplemental Figure S5** and **S6**). Other important features were low gene density and enrichment for PRC1 (Ring1b, H2AK119ub1, Rybp) and PRC2 (Ezh2) at (fast) silenced genes in both models, with PRC1/2 playing a more prominent role in the silencing dynamics model (MDA for Ring1B of 5% in silencing dynamics and 3% in XCI/escape model).

Among the top features specific for the XCI/escape model (Figure 4) we found the association with O-GlcNAc transferase Ogt (MDA 3-5%), which regulates a series of chromatin modifiers (e.g. Ezh2) and transcriptional regulators (e.g. RNA Pol II, Hcf1) (Yang and Qian 2017). Top features associated with silenced genes include the histone deacetylase HDAC1, involved in gene repression, as well as several features associated with active transcription, such as the H3K4 methyltransferase Mll2 (MDA 10% for escape class), the positive transcriptional elongation complex b (P-TEFb) component Cdk9, E2F1 and Hcfc1, and the transcription factor cMyc (MDA ranging from 2 to 4%). The dynamics of silencing by contrast seem to be strongly influenced by the MafK transcription factor, which is enriched at slowly silenced genes (Figure 4b). Interestingly, several features related to 3D chromosome organisation also appear to be associated with distinct silencing dynamics: While genes located in close proximity to a TAD border tend to be silenced slow, genes tend to be silenced faster when they are close to a LAD or highly connected to other genomic regions based on HiC/HiCap data. In summary, we have identified different feature sets that appear to influence whether or not a gene is subject to XCI and also, whether silencing occurs with slow or fast dynamics.

### Forest-guided clustering of X-linked genes uncovers combinatorial rules of gene silencing

The variable importance analysis described above pinpoints the individual contribution of each feature to the classification problem, but does not identify combinations of features associated with different silencing pathways, which ultimately determine the silencing class of each gene. Moreover, a large number of features appear to be of similar importance (MDA 2-4%), making it difficult to prioritize the important factors in both models.

To enable a better interpretation of the results and identify the main rule sets within the complex combinatorial classification of the RF models, we implemented a forest-guided clustering approach to stratify the genes into subgroups according to different combinations of rules. This approach uses the proximity of each gene to all other genes within the Random Forest model to group genes that are regulated by the same set of genomic and epigenetic features (see Methods section). The number of groups is chosen such that each cluster has a low degree of class mixture (containing mainly genes from one class and none or only few genes from the other class) while maintaining a small number of clusters in total (see **Supplemental Figure S7** and **Supplemental Text S2**). The results are visualized in a heatmap (Figure 5a and Figure 6a) showing the genes (columns), grouped by cluster, and a subset of features (rows) selected based on whether they were significantly different across clusters (top 10 features, sorted by p-value, from an ANOVA test). A cluster stability analysis showed that the clusters of the silencing dynamics are very stable, while the clusters of the XCI/escape model model are more variable but no cluster dissolves (**Supplemental Figure S7** and **Supplemental Text S2** for further information).

**Figure 5:**
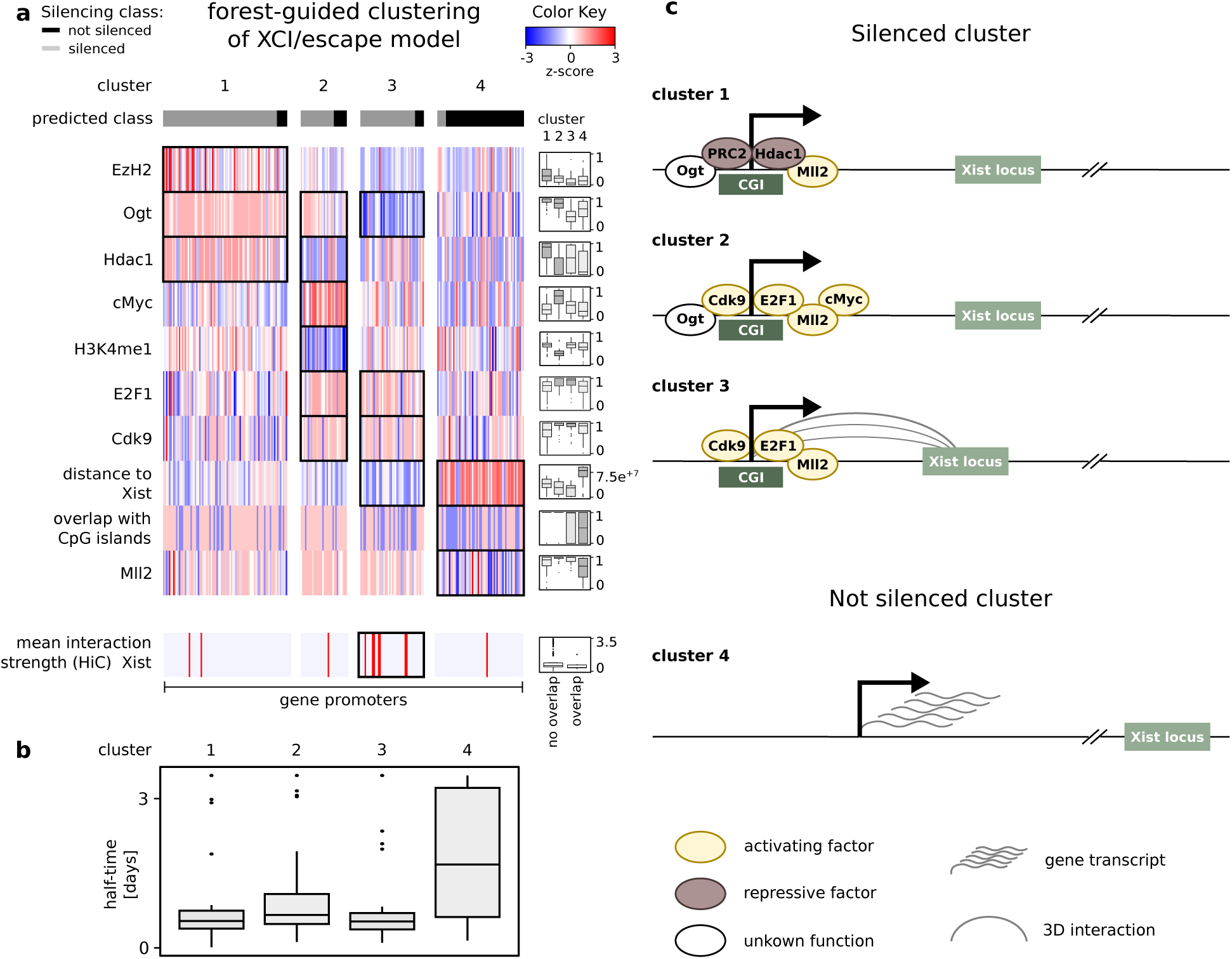
Classification rules for silenced vs not silenced genes derived from the XCI/escape model. (**a**) Forest-guided clustering reveals four main clusters of genes: elements in cluster 1, 2 and 3 mainly correspond to genes predicted from the model as “silenced” (marked in grey), while cluster 4 mainly contains genes predicted as “not silenced” (marked in black). Rows of the heatmap correspond to the top 10 features which showed significant differences among clusters, according to a p-value of an ANOVA test. Feature signals are scaled between −3 and 3 in each row separately, to allow a visual comparison of features from different scales. Scaled values are represented on a red-blue scale, with enriched features mainly displayed in red, and depleted features in blue. Differences in the distributions of features across different clusters are highlighted in the box plots next to the heatmap. (**b**) Distribution of the gene half-times estimated from the PRO-seq time series data for each cluster. (**c**) Schematic view of the distinct molecular mechanisms leading to gene silencing (cluster 1,2 and 3) or gene escape (cluster 4).

**Figure 6:**
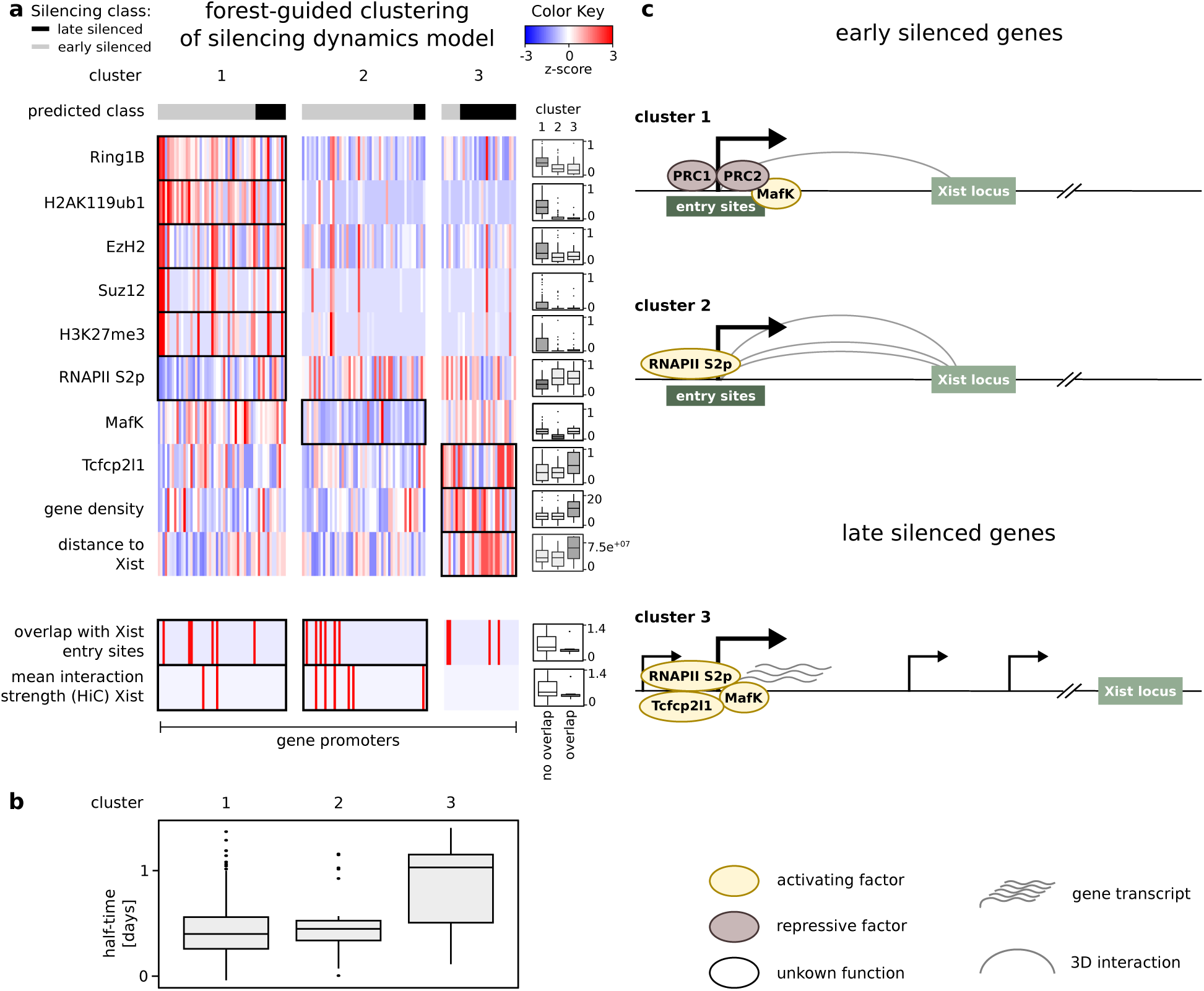
Classification rules for early vs late silenced genes derived from the silencing dynamics model. (**a**) Forest-guided clustering reveals three main clusters of genes: elements in cluster 1 and 2 mainly correspond to genes predicted from the model as “early silenced” (marked in grey), while cluster 3 mainly contains genes predicted as “late silenced” (marked in black). Rows of the heatmap correspond to the top 10 features which showed significant differences among clusters, according to a p-value of an ANOVA test. Feature signals are scaled between −3 and 3 in each row separately, to allow a visual comparison of features from different scales. Scaled values are represented on a red-blue scale, with enriched features mainly displayed in red, and depleted features in blue. Differences in the distributions of features across different clusters are highlighted in the box plots next to the heatmap. (**b**) Distribution of the gene half-times estimated from the PRO-seq time series data for each cluster. (**c**) Schematic view of the distinct molecular mechanisms leading to fast gene silencing (cluster 1,2) or slow gene silencing (cluster 3).

For the XCI/escape model four clusters are found (Figure 5a). Clusters 1, 2 and 3 are mainly populated with genes predicted as silenced, while cluster 4 contains not silenced genes (Figure 5b). Generally, genes tend to escape silencing when they are far from the *Xist* locus, when they have low levels of Mll2 at their promoters and when they do not have a CpG island (cluster 4). A large cluster of silenced genes (cluster 1) is already marked by a repressive chromatin state (PRC1/2, Hdac1) (**Supplemental Figure S8** and **S9**), while the two smaller clusters are bound by E2F1 and Cdk9, which are associated with active transcription (cluster 2 and 3). In addition, cluster 1 and 2 are enriched for OGT, cluster 2 is also enriched for binding of cMyc and genes in cluster 3 are particularly close to the *Xist* locus (Figure 5a and **Supplemental Figure S8**). Taken together, genes pre-marked by PRC1 and 2 as well as Hdac are generally silenced, while genes at the distal ends of the X chromosome tend to escape silencing. Moreover, genes associated with features of active transcription will be silenced, if they are enriched for Ogt and cMyc (cluster 2) or if they are very close to the Xic with strong 3D interactions with the Xist locus (cluster 3) (Figure 5c).

In the next step, in the silencing dynamics model, we focused on genes that become silenced by Xist and investigated the factors that would distinguish fast and slowly silenced genes. Here, the forest-guided clustering approach produced three clusters (Figure 6a): clusters 1 and 2 contain genes with lower half-times and are therefore predicted as early silenced, and cluster 3 is enriched for genes with higher half-times and predicted as late silenced (Figure 6b). Gene promoters in both early silenced clusters (1 and 2) tend to be close to the Xist locus (Figure 6a and Figure 6c), similar to the silenced genes in the analysis above (Figure 5a). Again, a fast silenced cluster (1), pre-marked by polycomb-repressed chromatin (H2AK119ub1, Ring1B, EzH2, Suz12, H3K27me3) is found. The second fast silenced cluster is mainly characterized by depletion of the transcription factor MafK and moderate enrichment of features related to transcriptional elongation, such as Ser2-phosphorylated RNA Polymerase II and H3K36me3 (Figure 6a, **Supplemental Figure S10** and **S11**). Interestingly, genes overlapping with Xist entry sites and genes that exhibit 3D contacts with the *Xist* locus are preferentially found in the early silenced cluster 2 (Figure 6a bottom). The late silenced genes fall in cluster 3 and mainly differ from early silenced genes by their distance to the *Xist* locus and by their genomic context, with being preferentially located in gene dense regions. In addition, slowly silenced genes show a moderate enrichment of the transcription factor Tcfcp2l1. By taking the data from Sakata et al. (Sakata et al. 2017), we also analysed genes that can be silenced by an Xist mutant lacking the A repeat in the trophoblast, and found them to be enriched in the fast cluster 1 compared to the slow cluster 3 (p-value=0.07 Fisher exact test, **Supplemental Figure S12**). These genes probably rely on the polycomb pathway for silencing in the trophoblast and appear to the be pre-marked by polycomb already in undifferentiated ES cells. In summary, our clustering analysis reveals that PRC1/PRC2 pre-marking seems to promote rapid silencing, while a long genomic distance from the *Xist* locus will delay silencing. For genes that are close to Xist, but depleted for the PRC1/2-associated chromatin marks, a high connectivity within 3D chromatin organisation appears to accelerate silencing.

### Experimental testing of model predictions

To validate our machine learning model we used the trained XCI/Escape RF model to predict the silencing class for X-chromosomal genes that could not be analyzed in the PRO-seq experiment due to insufficient coverage (Figure 7a and **Supplemental Table S1**). Although the number of X-linked genes that are not included in our PRO-seq data set, but had polymorphisms and adequate expression levels prior to XCI were limited, we nevertheless identified two genes for assessment by pyrosequencing as validation of our models. The silencing dynamics of one gene, *Sat1*, predicted to be inactivated and another gene, *Wdr13*, predicted to escape, that had been selected based on prediction probability and expression level, were then assessed experimentally (Figure 7a). To this end, TX1072 cells were treated with doxycycline for 8h and the ratio of *Sat1* and *Wdr13* mRNA originating from the B6 and Cast chromosomes was quantified by pyrosequencing at different time points (Figure 7b). *Sat1* expression from the B6 chromosome was reduced by 50% after 8h doxycycline treatment and was clearly silenced more strongly than *Wdr13*, which stayed approximately constant during the time course (Figure 7b). We also estimated the silencing half-times for *Sat1* and *Wdr13* from a pyrosequencing experiment performed on RNA extracted from the same cells used for the initial PRO-seq experiments. As predicted, *Sat1* falls in the silenced class with half-time of 0.88 days, while *Wdr13* falls in the not silenced class with a half-time of 1.65 days (**Supplemental Figure S13**). These results confirm that our machine learning model can predict X-chromosomal gene silencing based solely on epigenetic and genomic features.

**Figure 7:**
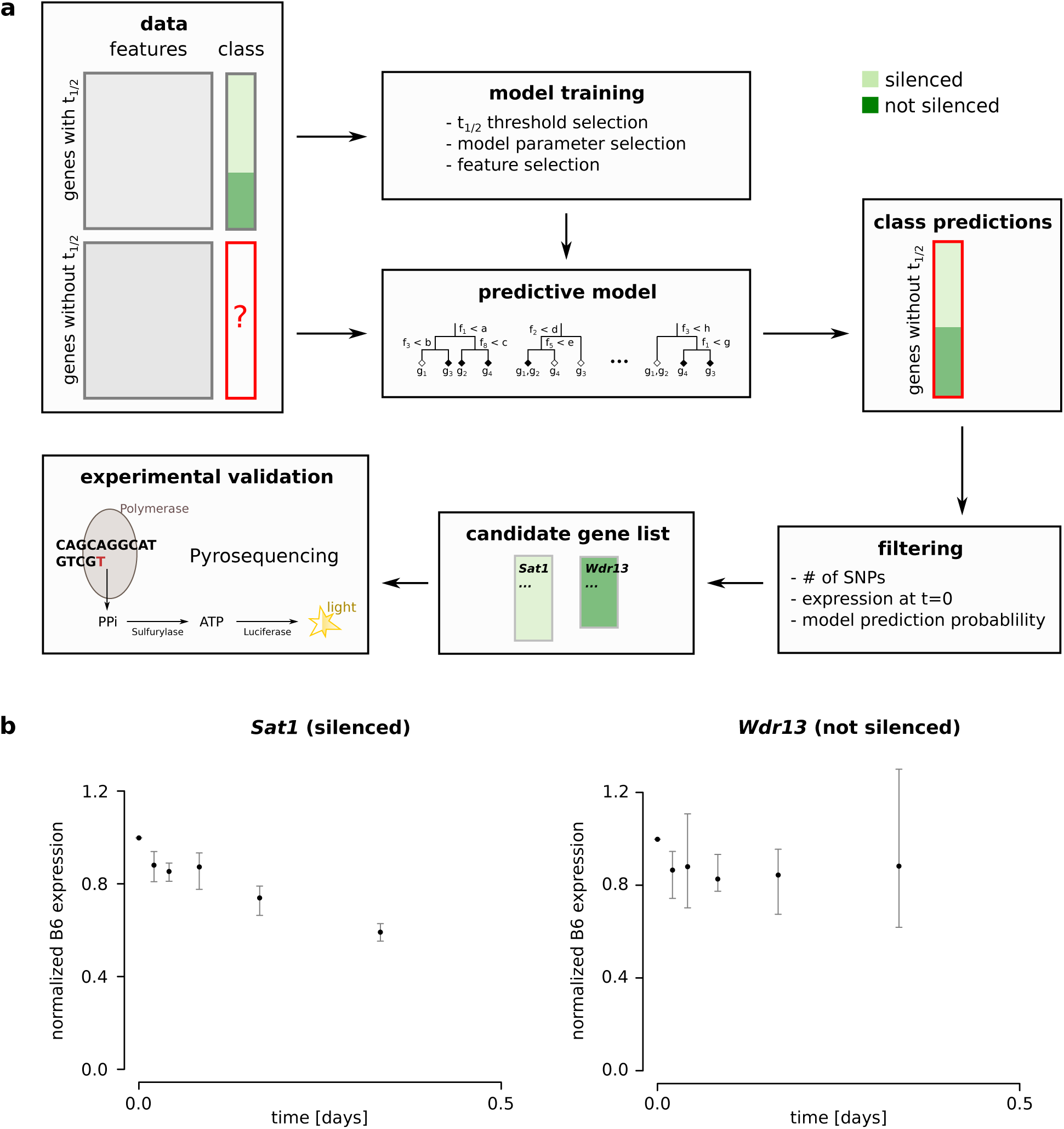
Chromosome-wide prediction of X-linked gene silencing and experimental validation of few candidates. (**a**) Workflow of gene silencing prediction and candidate selection for experiments. The XCI/escape model is trained on labeled data (gene promoter features and corresponding half-time class). The trained model is then used to predict the silencing class of all unlabeled X-linked genes, i.e. genes for which a half-time could not be estimated from the data, given the same set input features. Few candidate genes with newly predicted silencing class are chosen for experimental validation based on different filtering criteria. (**b**) Experimental validation of one gene predicted as silenced (*Sat1*) and another gene predicted as not silenced (*Wdr13*). Allele-specific quantification of *Sat1* (left) and *Wdr13* (right) through pyrosequencing at different time points during 8h of doxycyline treatment in TX1072 cells. The measurements are corrected for basal skewing and normalized to the uninduced control (for details see methods). Mean and standard deviation of 3 replicates are shown.

## Discussion

In this study we have developed a machine learning model that can predict a genes susceptibility to Xist-mediated silencing from a combination of epigenetic and genomic features. To train the model we measured silencing kinetics with high temporal resolution through allele-specific PRO-seq. Compared to previous studies (Marks et al. 2015; Borensztein et al. 2017), we assess silencing dynamics by measuring nascent transcription, therefore observing instantaneous changes in transcription by transcriptionally engaged PolII. Such measures of “transcription turn off” are not complicated by preexisting levels of mRNAs, as well as RNA processing and degradation dynamics. Moreover, the use of an inducible system allows us, in contrast to a previous study (Marks et al. 2015), to uncouple XCI from differentiation and to avoid the use of mutations in Xist cis-regulation to ensure non-random XCI. To identify the rules that govern the silencing dynamics, we developed a statistical random-forest model that can predict gene silencing dynamics from a set of genomic and epigenetic features. Unlike previous studies that focus on just a few sets of genes and/or investigate a few selected promoter features which are potentially linked to the XCI (Kelsey et al. 2015; Marks et al. 2015; Loda et al. 2017), we set out to identify silencing determinants in an unbiased manner based on a large number of epigenetic and genomic features. In order to discern the combinatorial rules which play a role in silencing dynamics we go one step beyond classical variable importance analysis in Random Forests and introduce a visualization scheme, based on Random-Forest-based clustering analysis. The determinants of silencing for groups of clustered genes from both the XCI/escape model and the silencing dynamics model retrieve previous observations but also shed light on novel players or combination of features which seem to have an important role in a genes susceptibility to Xist-mediated inactivation.

At the onset of XCI, Xist RNA does not cover the whole X chromosome evenly, but probably spreads by proximity transfer to regions that exhibit 3D contacts with the *Xist* locus (Engreitz et al. 2013). Contact frequency, and consequently also Xist coating, generally correlates with the genomic distance from the *Xist* locus, but is enriched at a set of 28 “Xist entry sites” that are spread along the X chromosome (Engreitz et al. 2013). In agreement with these observations our model identifies the genomic distance to the *Xist* locus as the primary determinant of gene silencing dynamics, an association that was also described previously (Marks et al. 2015). Moreover, we find that genes with strong 3D interactions with the *Xist* locus and genes that overlap with Xist entry sites are generally silenced fast, which has also been seen during imprinted XCI in preimplantation embryos (Borensztein et al. 2017). Interestingly, Xist initially tends to spread to gene-dense regions (Engreitz et al. 2013; Simon et al. 2013), but in our analysis, gene density is associated with reduced silencing, suggesting that Xist coating is not the only determinant of silencing.

Upon coating of the X chromosome, Xist recruits several protein complexes that then mediate gene silencing. While the protein Spen has been shown to directly bind to the A-repeat on the Xist RNA and be required for silencing in ES cells (Wutz et al. 2002; Chu et al. 2015; Monfort et al. 2015), a subset of genes in the extraembryonic trophoblast, where XCI is imprinted, can be silenced by an Xist mutant lacking the A-repeat (Sakata et al. 2017). Those genes might be particularly sensitive to alternative, Spen-independent silencing pathways, potentially mediated by the polycomb repressive complexes, which have been shown to play a more prominent role for silencing in the trophoblast compared to ES cells (Brockdorff 2017). Interestingly, these A-repeat independent genes are silenced particularly fast in our data set and are enriched in a cluster of rapidly silenced genes (Figure 6a) that is enriched for PRC1 and to a lesser extend PRC2. This finding suggests that polycomb pre-marking prior to the onset of XCI might accelerate gene silencing, either by enhancing recruitment of additional repressive complexes or by helping to recruit Xist to gene promoters. A similar enrichment of PRC2 chromatin mark H3K27me3 and the PRC1 component Ring1b has previously been found at genes susceptible to ectopic silencing by Xist transgenes (Kelsey et al. 2015; Loda et al. 2017).

In addition to pre-marking with repressive chromatin modification and complexes (PRC1/2, Hdac), we surprisingly also find features associated with active transcription at promoters of silenced genes. The H3K4 methyltransferase Mll2/Kmt2b, which deposits H3K4me3 at bivalent promoters in ES cells (Hu et al. 2013; Denissov et al. 2014), and also Hcfc1 that is part of the Mll2 complex (Herz et al. 2013) and E2F1 that recruits the Mll2 complex in a cell cycle dependent manner (Tyagi et al. 2007), are enriched at silenced genes. Moreover, the transcription factor cMyc and the Cdk9 protein, a member of P-TEFb, are found at a subset of silenced genes. Interestingly, a large number of silenced genes are strongly enriched for Ogt, the single enzyme that catalyzes the post-translational modification O-GlcNAc, found at Ser/Thr residues in many proteins (Yang and Qian 2017). For example Hcfc1 activity requires O-GlcNAcylation and the PRC2 component Ezh2 is stabilized by O/GlcNAcylation (Yang and Qian 2017). Moreover, RNA Pol II is O-GlcNAcylated at Ser2 and Ser5, thus competing with phosphorylation (by Cdk9 for Ser2) of these residues, which is required for transcription initiation (Ranuncolo et al. 2012; Lewis et al. 2016; Harlen and Churchman 2017). Since Ogt targets a large number of proteins and affects transcription in many different ways, it is hard to pinpoint at this moment on how it might promote Xist-mediated silencing, but our finding opens up an interesting avenue for future studies. It should also be noted that the Ogt gene is actually X-linked, raising the intriguing possibility that it might even be implicated in the XX dosage sensitivity involved in XCI.

Finally, our analysis identified several structural features that appear to modulate the dynamics of silencing. A high “connectivity” of some genes, i.e. how much the gene is involved in 3D interactions with other genomic elements, is associated with faster silencing, maybe because Xist RNA can spread more easily to these genes through proximity transfer. Moreover, fast silencing preferentially occurs at genes that are close to a LAD, which generally contain repressed genes (van Steensel and Belmont 2017), while genes close to TAD boundaries tend to be silenced slowly.

In conclusion, we have developed two Random Forest models that can accurately predict silenced and not silenced/escape genes, but also classes of early versus late silenced genes, constituting the first chromosome-wide predictive models of gene silencing from a very large set of features. We confirmed the predictive nature of our models by experimental testing of model predictions. The Random Forest approach allows us to quantify the relative contribution of several features that have previously been associated with XCI (e.g. linear distance to *Xist*, enrichment for PRC1 and PRC2 etc) and suggested new features that can be tested in more detail in future studies. It is however likely that additional features, which are not included in the current model due to missing or poor quality datasets, might contribute to modulate the susceptibility to Xist-mediated silencing. For example, certain repetitive elements, such as LINEs have been suggested to affect silencing and escape from XCI (Lyon 1998). Including additional features in the future will likely further improve our ability to predict silencing susceptibility and a detailed experimental investigation of the different silencing pathways elicited by Xist will facilitate the interpretation of the features that predict silencing dynamics as well as escape from XCI.

## Methods

### ES cell culture

The female TX1072 cell line is a F1 hybrid ESC line derived from a cross between the 57BL/6 (B6) and CAST/EiJ (Cast) mouse strains that carries a doxycycline responsive promoter in front of the *Xist* gene on the B6 chromosome and an rtTA insertion in the *Rosa26* locus (described in Schulz et al. (Schulz et al. 2014)). TX1072 cells were grown on gelatin-coated flasks in serum-containing ES cell medium (DMEM (Sigma), 15% FBS (Gibco), 0.1mM β-mercaptoethanol, 1000 U/ml leukemia inhibitory factor (LIF, Chemicon)), supplemented with 2i (3 μM Gsk3 inhibitor CT-99021, 1 μM MEK inhibitor PD0325901). Cells were seeded at a density of 10^5^ cells/cm^2^ coated with gelatin two days before the experiment. Xist was induced by supplementing the medium with 1 μg/ml Doxycycline. For PRO-seq samples were collected before doxycycline treatment (0h) and at time points 0.5, 1, 2, 4, 8, 12 and 24 h after treatment. Samples without doxycycline and 24 h doxycycline were collected in duplicate. For mRNA-Seq on undifferentiated TX1072 cells, samples were collected before doxycycline treatment and at time points 2, 4, 8, 12 and 24 h after treatment. Also for mRNA-seq samples without doxycycline and 24 h doxycycline were collected in duplicate. For mRNA-Seq on differentiating TX1072 cells samples were collected at time points 0, 8, 16, 24 and 48h.

### PRO-Seq

For each timepoint ∼1 × 10^7^ nuclei were isolated by washing the cells twice with ice-cold PBS, and once with 15 ml swelling buffer (10 mM Tris-Cl, pH 7.4, 300 mM Sucrose, 3 mM CaCl_2_, 2 mM MgAc_2_, 5 mM DTT). Then, 15 ml cell lysis buffer (10 mM Tris-Cl, pH 7.4, 300 mM Sucrose, 3 mM CaCl_2_, 2 mM MgAc_2_, 0.5% NP-40, 1 mM PMSF, EDTA-free protease inhibitors (1 tablet for 50 ml buffer; Roche), 5 mM DTT) is added and cells are scraped off the plate into a 50 ml tube and spun at 900 g and 4 ^°^C in a swing bucket centrifuge for 5 minutes. Supernatant is removed and the cell pellet is resuspended in 5 ml cell lysis buffer, transferred to a 7 ml dounce homogenizer and dounced 50x on ice. Dounced cells are moved to 15 ml tube and spun at 1200 g and 4 ^°^C in a swing bucket centrifuge for 5 minutes. Supernatant is removed and the nuclei are counted, snap frozen and stored in glycerol storage buffer (50 mM Tris-Cl, pH 8.3, 40% glycerol, 0.1 mM EDTA, 5 mM MgAc_2_, 1 mM PMSF, EDTA-free protease inhibitors (1 tablet for 50 ml buffer; Roche), 5 mM DTT).

Run-on and library preparation was performed as previously described (Mahat et al. 2016), using the single biotin-CTP nucleotide run-on protocol to prolong run-on and increase sequence length. In short, run-on was performed with 1 × 10^7^ nuclei in 100 ml glycerol storage buffer and 100 ml pre-heated nuclear run-on mix, to get a final concentration in the run-on of 5 mM Tris-HCl, pH 8, 2.5 mM MgCl2, 0.5 mM DTT, 150 mM KCl, 0.025 mM biotin-11-CTP, 0.25 mM CTP, 0.125 mM ATP, UTP and GTP, 0.5% sarkosyl and RNase inhibitor. Run-on was done for 5 minutes at 37 ^°^C and stopped by adding 500 ml Trizol LS. RNA isolation, base hydrolysis, biotinylated-RNA enrichment steps, enzymatic modifications of RNA, adapter ligations, reverse transcription, amplification and library size selection were done as described previously(Mahat et al. 2016). Libraries were sequenced on the HiSeq 2000 Illumina sequencer (100bp, single-end). For each library at least 50 Mio reads were generated.

### Processing of PRO-Seq data

Adapter sequences were trimmed with cutadapt v1.8.2. Nucleotides with poor 3’ base quality (BAPQ < 20) were trimmed and reads of <20 bp were discarded. After quality control between 30 to 50 million reads remained. Ribosomal reads were first removed by alignment to the rRNA reference (GenBank identifiers:18S, NR_003278.3; 28S, NR_003279.1; 5S, D14832.1; and 5.8S, KO1367.1) using bowtie1 (v1.0.0) and allowing 2 mismatches in the seed (-m 1 -l 20 -n 2 options). Then, non-ribosomal reads were mapped to both parental genomes. To do this, the VCF file (mgp.v5.merged.snps_all.dbSNP142.vcf) reporting all SNP sites from 36 mouse strains, based on mm10, was downloaded from the Sanger database. SNPsplit tool (v0.3.0) was used to reconstruct the Cast genome from the mm10 reference. Only random best alignments with fewer than two mismatches (-M 1 -v 2 -l 20 options) were kept for downstream analyses. We applied an allele-specific RNA-seq strategy as described in Borensztein et al.(Borensztein et al. 2017). Briefly, mapping files of both parental genomes were merged for each sample and SAMtools mpileup (v1.1) was then used to extract the base-pair information at each genomic position. Read counts mapping to the paternal and maternal genomes, respectively, were summed up across all SNPs present in the same gene. To avoid allele specific bias, we checked the genotypes using a ChIP-Seq input from the same cell line (data not shown). Therefore, only SNPs covered by at least 10 reads in this input sample and having an allelic ratio range between 0.25 and 0.75 were kept for downstream analysis (17,035,327 SNPs in total). RPKM values were calculated using gene count table, generated with Gencode annotation (M9) and HTSeq software (v0.6.1).

### RNA extraction and cDNA preparation

Cells were lysed by direct addition of 1 ml Trizol (Invitrogen). Then 200μl of Chloroform was added and after 15 min centrifugation (12000xg, 4°C) the aqueous phase was mixed with 700 μl 70% ethanol and applied to a Silica column (Qiagen RNAeasy Mini kit). RNA was then purified according to the manufacturers recommendations, including on-column DNAse digestion. Concentration and purity were checked on a Nanodrop. In case of a low 260/230 ratio, extra ethanol precipitation was performed. RNA profiles were then checked by Bioanalyzer (Agilent RNA 6000 Nano kit) and 1ug of RNA from each condition was used for mRNA-seq. For pyrosequencing, 1ug RNA was reverse transcribed into cDNA using Superscript III Reverse Transcriptase (Invitrogen).

### Pyrosequencing

For allele-specific expression analysis of *Sat1* and *Wdr13*, pyrosequencing technology was used. Starting from cDNA, an amplicon containing a SNP is amplified by PCR with primers CGACACTTCATGGCAACCTAGTA, AAGAGGGTGAAATGTTCTCTCTGG (reverse) for *Sat1* and GTCAACTCTGCCACCTCAAAAATT, GCAACAGAATTTGGGTACATAACA for *Wdr13* using GoTag Flexi G2 (Promega) with 2.5 mM MgCl2 for 40 cycles. The PCR product was sequenced using the Pyromark Q24 system (Qiagen) with primers CTTCATGGCAACCTAGTA for *Sat1* and TTCATCACCAATCATCC for *Wdr13*.

### mRNA-Seq

The 32 libraries were prepared with an input of 1µg of totalRNA, using the *TruSeq Stranded mRNA LT Sample Prep kit* from Illumina, according to manufacturer’s recommendations. Single Index kit was used, and 12 cycles of PCR were set up. Final libraries were quantified with *Qubit dsDNA HS Assay Kit*, and qualified with *LabChip® GX system* (PerkinElmer). Then 2 equimolar pools of 16 libraries each were prepared at 10nM. The exact molarity of the pools were assess by qPCR using the *KAPA Library Quantification Kit Illumina* on CFX96 system (Biorad). Then each pool was sequenced on 1 flowcell of HiSeq 2000 system (paired-end, 100bp reads) in PE100, in order to target ∼100M cluster per sample.

### Processing of mRNA-Seq data

First ten bases from all reads were removed, due to their low quality, using FASTX toolkit (v0.0.13). Reads were then mapped to both parental genomes with TopHat2 software (v2.1.0). Only random best alignments with less than two mismatches were kept for downstream analyses. We applied the same allele-specific RNA-seq strategy used for PROseq data analysis.

### Silencing half-times

From the allele-specific counts, the fraction of reads mapping to the B6 chromosome were calculated for each gene as the reads mapped on B6 (mm10) genome divided by the total number of allele-specific reads.

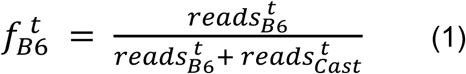

Only genes with a minimum of 10 allele-specific reads at each time point were considered for further analysis. 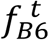 was averaged across replicates (0, 24h), resulting in a total of eight time points (t = 0, 0.5, 1, 2, 4, 8, 12, 24h). For estimating gene-specific silencing half-times we first normalized the data to the uninduced control and corrected for basal skewing (different transcriptional activity at the two alleles in the absence of dox). We assume that transcription is constant on the Cast allele throughout the time course since Xist is induced only on the B6 chromosome. Solving (1) for 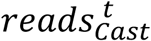 at t=0h gives

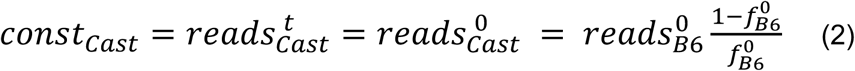

If we solve (1) for 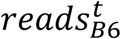 and substitute 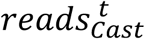 with (2) we obtain:

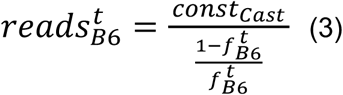

we can calculate the expression from the B6 allele relative to the uninduced control (t=0) 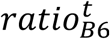 for each time point as follows

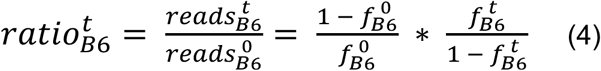

Silencing half-times were estimated by fitting *ratio* 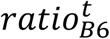 with an exponential decay function:

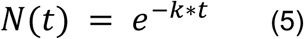

where *k* represents the silencing rate and *N*(*t*) the fit for 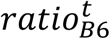. The nonlinear least-squares estimate of the parameter *k* of the exponential decay function in (5) was determine with the `nls` function (`stats` R package). For each gene the half-time, defined as the time point at which gene expression is reduced to 50% of its initial value at *t=0* is estimated from *k*:

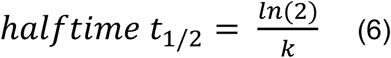

A maximum value of *k* = 5, corresponding to a half-time of 3.5 days was set, as higher half-times cannot be reliably estimated from our data, due to the limited range of time points from 0 to 24 h. The goodness of fit was evaluated via the square root of the residual sum of square *sqrtRSS* defined as:

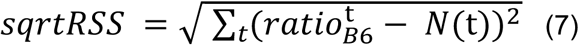

In order to obtain reliable half-times, only genes with a *sqrtRSS* smaller than 1.5 and a basal skewing 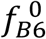 between 0.2 and 0.8 were considered for further analysis. From the PRO-seq data, we could compute half-times for 296 X chromosomal genes on mouse genome mm10 (**Supplemental Table S4** and **Supplemental Table S2**). Those genes were mapped to the mouse genome mm9 with the `liftOver` tool (Kuhn et al. 2007) from UCSC Genome Browser. Gene half-times from both differentiated and undifferentiated mRNA-seq time series were computed in the same way as described above. For the undifferentiated mRNA-Seq data set we discarded replicate B due to insufficient read coverage and only used replicate A (Figure 2a), which resulted in computing half-times for 346 genes. For the differentiated mRNA-Seq data set we averaged replicate A and B for each time point and computed half-times for 379 genes. For 233 genes, half-time could be estimated from all 3 data sets.

### Identification of active gene TSSs

To identify for each gene the transcription start site (TSS) that is used in embryonic stem cells, we annotated regulatory regions (RR) based on the PRO-seq data with the dREG method (Danko et al. 2015). RRs are defined as regions which harbor bidirectional transcription from the PRO-seq signal at time point t=0. Both replicates at t=0 were analysed separately and de-novo RRs with a quality score of 0.8 or higher were selected. Those RRs are indicative of active transcription start sites (TSS) and were used to assign each gene to its active TSS (**Supplemental Figure S3**). Regulatory regions with overlapping genomic ranges between replicates were merged into one region. Most of the identified RRs overlapped known gene promoters. If an RR was found within +/- 100bp of an annotated gene TSS, the TSS was chosen as active TSS for that gene. If multiple gene TSSs were found to overlap RRs, the active TSS overlapping the RR with the strongest signal (i.e. highest score) was chosen for that gene. If no RR was found within +/- 100bp of an annotated gene TSS, the genomic search space was extended to +/- 1000bp. If an RR could be found within +/- 1000bp of an annotated gene TSS, a novel alternative TSS, coincident with the middle point of the RR, was defined for that gene. If no RR could be found also within the enlarged region, the gene was discarded. This revised gene TSS annotation for 280 genes with computed half-times was used to quantify promoter features from ChiP-Seq data sets (**Supplemental Table S4**).

### Definition of model features

The epigenetic and genomic features used for the Random Forest models are listed in detail in **Supplemental Table S3** and summarized in Table 1. In total, 133 ChIP-seq libraries and one bisulfite sequencing experiment on undifferentiated mouse embryonic stem cells (mESC) were collected from various sources. After performing ChIP-Seq library quality control with the *deepTools* package (Ramírez et al. 2014) and stringent filtering, 57 ChIP-seq libraries and the bisulfite sequencing experiment were used in the model (**Supplemental Text S1** for details on data pre-processing and filtering and **Supplemental Figure S14** and **S15**). Epigenetic features are defined as the average ChIP-seq signal in a pre-defined region around the genes TSS, normalized to the signal of a matched control experiment in the same region (see **Supplemental Figure S16** as example). Read counts of each feature were normalized to the control with the R package `normR` (Helmuth et al. 2016; Kinkley et al. 2016). The pre-defined regions from which the epigenetic signal is computed are defined individually for each feature to account for broad and narrow features, e.g. epigenetic signal for H3K36me3 vs. Transcription Factor signal cMyc, by manually inspecting the heatmap plots generated by the deepTools (**Supplemental Figure S14** and **S15**, **Supplemental Table S3).**

In addition to epigenetic features, we defined 16 genomic features, including distance of each genes TSS to the Xist locus or to the next TAD border. A gene was considered to overlap with a LAD if a region of 1000 bp around its TSS overlapped with an annotated LAD. Gene density was defined as the number of annotated genes within the 200 kb region around the TSS. 3D interactions were quantified as the number of interactions (number of loci that interact with the gene) or strength of interactions (average read counts / number of interactions) defined by HiC or HiCap data for each genes promoter. A gene was classified as overlapping with a CpG island if a region of 1000 bp around each genes TSS overlaps with an CpG island as annotated in the UCSC genome browser (mm9). CpG content was defined as the normalized CpG content within the 1000bp region around each genes TSS, computed as the ratio of observed over expected CG dinucleotides (Marsico et al. 2013). A detailed description of all genomic features can be found in **Supplemental Text S1**.

### Random Forest classification models

Two statistical models were developed to predict “silenced” vs “not silenced” genes, referred to as “XCI/escape model” and the other one to distinguish “early” vs “late” silenced gene, referred to as “silencing dynamics model”. The continuous half-time values were therefore assigned to discrete classes in both models, according to fixed thresholds, which were chosen such that the error rate from the classification model (see below) would be minimized (Figure 1g, Figure 2g and **Supplemental Table S6**). Genes were defined as silenced for *t*_*1/2*_ *< 0.9* and as not silenced for *t*_*1/2*_ *> 1.6*; they were classified as “early silenced” if *t*_*1/2*_ *< 0.6* and as “late silenced” if 0.9 < *t*_*1/2*_ *< 1.4* Genes outside these ranges were not included in model training. At the end, the XCI/escape model was trained on 218 genes (168 from the “silenced” and 50 from the “not silenced” class) and the silencing dynamics model on 139 genes (100 from the “early” silenced and 39 from the “late” silenced class).

The two Random Forest classification models were implemented with the `randomForest` R package. Random forests are non-parametric classifiers which make use of multiple decision trees to learn non-linear classification tasks. The use of multiple trees makes the method robust to outliers and noise, and reduces the risk of overfitting, also with a small number of training examples, strong class imbalance and correlated features. Class imbalance is present in both our data sets (168 silenced versus only 50 not silenced genes and 100 early silenced versus 39 late silenced genes), as well as correlation between epigenetic and/or genomic features (**Supplemental Figure S17**). In a Random Forest, for each tree a random subset of training genes are drawn with replacement from the whole dataset. We set this number (*sampsize* parameter in the `randomForest` R package) to the *size of the smaller class - 10* for both classes, to ensure that each tree is trained on a balanced subset of the data, thereby avoiding a classification in favour of the larger class. The examples of the dataset which are not used by a classification tree for training (out-of-bag data) constitute a test set for that particular tree and are used to compute the prediction error of the tree, the out-of-bag (OOB) error.

The prediction for each gene is made by taking a majority vote from the predictions over all trees for which that sample was part of the out-of-bag data. By comparing the OOB predictions with the measured half-time training set one can estimate the prediction error rate. We used Random Forest classification with 1000 trees to predict the silencing class for our X-chromosomal genes in both classification settings (i.e. XCI/escape model and silencing dynamics model) using in total 74 predictor variables (epigenetic and genomic features). The *mtry* parameter, defining how many features are randomly tested at each split in the tree, was optimized during training such that the OOB error of the corresponding Random Forest is minimized (i.e. the average OOB error from all the trees).

Random Forest provides several internal measures of feature importance, based on out-of-bag data. Here, we chose the *mean decrease in accuracy* (MDA) as feature importance criterion because it has a straightforward interpretation. The MDA for a given feature is the decrease in model accuracy from permuting the values of that feature, averaged over all trees. Therefore variables with large positive values of the MDA correspond to important features for the classification, while variables with MDA close to zero or negative correspond to unimportant features or noise. MDA is computed for every feature in the model and for each class separately, as some features might contribute more to the prediction of one class vs the other. Feature importance (MDA) and classification performance (OOB error) measures were further averaged over a collection of five hundred Random Forests to obtain stable results.

Simple feature selection was performed to improve the model performance by removing weaker or redundant features, which potentially introduce noise. We ranked features in each class according to their MDA and computed the models error rate on the top *x* features from both classes (including only those variables with MDA > 0). *x* was optimized to obtain the combination with minimal error rate. The *top feature* set with minimal error rate (10 from each class of the XCI/escape model and 11 from each class for the silencing dynamics model) was then used to train a second collection of Random Forests. For both models, The classification performance is reported as average of five hundred Random Forests trained on the *top features*.

### Forest-guided clustering for model interpretation

In order to unravel the combinatorial rules of X-chromosomal gene silencing, as well as correlations between important epigenetic and genomic features, we attempted to cluster and visualize X chromosomal genes according to those rules. Each individual tree in the Random Forest model contains several terminal nodes (i.e. leaves) with only a small number of observations (i.e. genes) which belong to one of the two classes. We can then extract a similarity measure between those observations: if two genes *i* and *j* land in the same terminal node, the similarity between i and j is increased by one(Breiman 2001). We computed similarities for all gene pairs and build an NxN symmetric matrix (with N=total number of genes), which we refer to as *proximity matrix* (Figure 3c). Each entry in the *proximity matrix* lies in the interval [0,1] and represents the frequency with which two genes occur in the same terminal node of a tree, intuitively defining “how close” two genes are in the forest. Next, the similarity values of this matrix are converted to dissimilarities or distances:

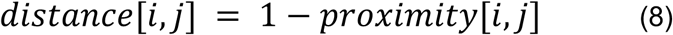

and used as input to k-medoids clustering(Reynolds et al. 2006) in order to group genes into clusters, using the `pam` function of the `cluster` R-package. The proximity matrix values and the class predictions used for clustering are also averaged over the 500 Random Forest models. As the clustering process is guided by the *proximity matrix* derived from the Random Forest, genes of the same silencing class (e.g. not silenced) are largely expected to cluster together according to a certain combination of epigenetic and genomic features. Given the non-linear nature of the classification problem modeled here, we also expect, to some extent, genes from the same silencing class to be grouped in different cluster according to different combination of features.

Similarly to k-means clustering, k-medoids clustering requires setting in advance the number of clusters *k*. We developed a scoring system to choose the optimal *k* which minimizes model bias and restricts model complexity (**Supplemental Text S2**). According to this scoring system, we chose k=4 for clustering genes from the XCI/escape model and k=3 for clustering genes from the silencing dynamics model (**Supplemental Figure S7**). The results of the k-medoids clustering are visualized for both models as heatmaps. As displaying all the models variables in the heatmaps would make visual interpretation harder, we display only the top 10 features which have a significant variation across clusters according to the p-value of an ANOVA test. Compared to classical Random Forest feature importance, the outcome of the forest-guided clustering enables an alternative interpretation of the Random Forest predictions in terms of combinatorial rules which determine the silencing state of groups of genes.

### Gene class predictions

Given our trained XCI/escape model we predicted the silencing class of all X-linked genes which were not included in the training set, either because of insufficient read coverage from the PRO-seq data or because of a poor fit to the exponential model. For these genes, we computed the same epigenetic features at gene promoters, as well as all genomic features as described above and gave them as input to the Random Forest model. After class prediction, few genes were chosen for experimental validation (Figure 7a) according to the following criteria: 1) sufficient expression for experimental detection at time point 0 (PRO-seq RPKM > 1, based on non-allele specific mapping); 2) at least one polymorphic site (SNP) in exonic regions and 3) probability of a gene to be predicted in a certain class (silenced vs not silenced) higher than 65%, averaged over 500 trained Random Forests.

### Statistical tests

A non-parametric Kolmogorov-Smirnov (KS) test was performed to test whether silencing dynamics significantly differ between A-repeat dependent genes and A-repeat independent genes. A Fisher exact test was performed to test whether known escapee genes (**Supplemental Table S2**) were enriched among: 1) genes with measured half-times higher than 1.6, which define the not silenced class in the XCI/escape model and 2) genes predicted as not silenced from the XCI/escape model. An Analysis of Variance (ANOVA) test was performed to test whether the distribution of the values of each features differed significantly across clusters in both the XCI/escape and the silencing dynamics model. All statistical tests were performed in R with the *base* statistical functions package.

## Data Access

The time-course RNA-seq data, for both differentiated and undifferentiated mESC, as well as the time-course PRO-seq data have been deposited in the Sequence Read Archive under accession code GSE (**pending for approval**). These data also include processed data which may serve for future analysis.

## Acknowledgements

We thank the NGS platform of the Curie Institute for library preparation, sequencing, data preparation and quality analysis reports, J. Helmuth for the help with ChIP-Seq data normalisation and Quentin Deletang for help with the mRNA-seq preparation.

This work was funded by NIH grant GM025332 to JTL, Center for Vertebrate Genomics (Cornell University, Ithaca, NY, USA) stipend to JTL and IJ, Helen Hay Whitney Postdoctoral Fellowship and Rosalind Franklin Fellowship awarded by the University of Groningen to IJ. E. Heard is supported by ERC Advanced Investigator award (ERC-2014-AdG no. 671027), Labelisation La Ligue, FRM (grant DEI20151234398), ANR DoseX 2017, Labex DEEP (ANR-11-LBX-0044), part of the IDEX Idex PSL (ANR-10-IDEX-0001-02 PSL), and ABS4NGS (ANR-11-BINF-0001). JC was supported by the ARC Foundation for Cancer Research (France) and the ATIP-AVENIR program (INSERM, CNRS, France). LBAS is supported by the International Max Planck Research School for Computational Biology and Scientific Computing.

## Author contributions

JTL and EH conceived the PRO-seq experiment and strategy; AM, EGS and EH conceived the main idea of the computational analysis; EGS and IJ performed the PRO-seq experiment and preprocessing of the data; EGS generated the mESC; AM, EGS and LBAS conducted the investigation; LS and CJC performed the preliminary analysis of PRO.seq and mRNA-seq data; LBAS and AM performed statistical analysis and data modeling; JC and CP generated the mRNA-seq data, BF performed the Pyrosequencing experiment. LBAS, AM and EGS wrote the original draft; IJ, JTL and EH reviewed and edited the paper; JTL and EH financed the study; JTL, EH and AM provided resources.

