## Supplementary for "Kinetics of Xist-induced gene silencing can be predicted from combinations of epigenetic and genomic features"

### Supplemental Information

1. Supplemental Text 1: Definition of epigenetic and genomic features
2. Supplemental Text 2: Clustering Details
3. Supplemental Figure 1: Correlation of PRO-Seq replicates
4. Supplemental Figure 2: Xist Tsix region allelic track.
5. Supplemental Figure 3: Example of assignment of an X-linked gene to its active TSS.
6. Supplemental Figure 4: Model performance.
7. Supplemental Figure 5: Feature importance for the XCI/Escape model.
8. Supplemental Figure 6: Feature importance for the silencing dynamics model.
9. Supplemental Figure 7: Cluster stability analysis for optimal number of *k* clusters.
10. Supplemental Figure 8: Enriched features from the XCI/escape model clustering.
11. Supplemental Figure 9: Top features from the XCI/escape Random forest.
12. Supplemental Figure 10: Enriched features from the silencing dynamics model clustering.
13. Supplemental Figure 11: Top features from the silencing dynamics Random forest.
14. Supplemental Figure 12: Comparison of clustering results from both models with classification of genes in repeat A-dependent and repeat A-independent from Sakata et al.[14].
15. Supplemental Figure 13: Experimental validation of Sat1 and Wdr13 with Pyrosequencing.
16. Supplemental Figure 14: ChIP-Seq library filtering with deepTools heatmap.
17. Supplemental Figure 15: Example on how to select the best ChIP-seq data set for a given factor.
18. Supplemental Figure 16: Example of ChIP-Seq signal normalization with normR.
19. Supplemental Figure 17: Feature correlation matrix.
20. Supplemental Table 1: List of Random Forest predictions for genes without half-times.
21. Supplemental Table 2: List of genes with computed half-times.
22. Supplemental Table 3: Metadata of Random forest features.
23. Supplemental Table 4: Filtering steps in the computation of half-times.

24. Supplemental Table 5: Filtering steps in the ChIP library pre-processing.
25. Supplemental Table 6: Ranges of half-times for choosing class thresholds.

### Supplemental Text 1: Definition of epigenetic and genomic features

#### Definition of epigenetic features

##### Pre-processing of ChIP-Seq libraries

A collection of 133 ChIP-Seq libraries with matching control libraries were downloaded from the Gene Expression Omnibus database (Edgar et al. 2002) (see **Supplementary Data Set 2**). ChIP-Seq and control reads (downloaded as sra or fastq files) were aligned to mm9 genome with `Bowtie2` (with number of mismatches = 1). Obtained sam files were converted into bam using `samtools`, only keeping alignments with MAPQ score higher than 10. Replicates were pooled for further analysis to obtain better coverage. All ChiP libraries containing less than three million uniquely mapped reads were removed from the collection, as the read coverage would be too sparse to infer robust signals. The `deepTools` package (Ramírez et al. 2014) was used for quality control of the ChIP-seq data: fingerprints, created with the `plotFingerprint` function and heatmap summary plots, created with `bamCoverage`, `computeMatrix` and `plotHeatmap` functions, were created for each ChiP library and the corresponding control library. Fingerprint plots produce a profile of cumulative read coverage from bins of specified size across the genome and allow to assess the signal-to-noise ratio in ChIP-Seq samples, i.e. whether there is sufficient enrichment of signals versus background. Heatmap plots allow to assess the average distribution of both ChIP-seq and control signal of interest around pre-specified regions (e.g. promoters) in the genome. For example, for a ChIP-seq dataset on H3K4me1 we expect an average signal enrichment (i.e. a peak) around the gene transcriptional start sites for the experiment but not for the control. ChIP libraries were filtered by manual inspection based on the enrichment of experiment over control input signal (see **Supplementary Figure 14** as example).

For some features more than one ChIP-Seq library was downloaded, when experiments from different labs were available in GEO, and the most 'high-quality' dataset for each feature was chosen based on the signal to noise ratio (deepTools heatmap) and cumulative distribution of the reads from input and experiment (deepTools fingerprint). For example, for the feature CTCF one out of three available ChIP libraries was selected, based on the fingerprint and heatmap summary plots of the ChIP libraries (see **Supplementary Figure 15**). After completion of all filtering steps, we defined regions of enrichment for each of the remaining 57 ChIP-Seq libraries and used the normalized read counts in the specified region as epigenetics features for the Random Forest model (see **Supplementary Table 2** and **Supplementary Data Set 1**).

#### Normalization of ChIP-Seq signals

The R package `normR` (Helmuth et al. 2016; Kinkley et al. 2016) was used to normalize each ChIP library to the corresponding control library in order to remove the background signal. normR jointly models ChIP and control reads over the whole genome with a binomial m-component mixture model where one component models the background noise and the remaining m-1 components model the signal. In our case only a two-component model is used: one component to account for the background and one component to account for the ChIP signal. The fitted background component allows to inspect the enrichment in a certain genomic region and is used to compare ChIP read counts for that region to the expected read counts under the fitted background component (see **Supplementary Figure 16**). This model can then be used to calculate a normalized enrichment for each region, where the fold change of ChIP vs control read counts of each region is regularized (windows with zero counts get zero enrichment) and standardized (to values between zero and one, where zero means no enrichment and one means 100% enrichment), making read counts comparable between different ChIP experiments.

#### Bisulfite-Seq data

We computed the DNA methylation level (*DNA methylation (BS-Seq)*) of each gene's promoter using the whole genome bisulfite sequencing data in mESC from Stadler et al. (Stadler et al. 2011). For each

C in a CG context, the total number of reads and the number of methylated reads is given from which the percentage of methylation (# methylated reads / # total reads) can be computed. We then computed the average methylation level over all CG sites within a 1000 bp region around each gene's TSS.

### Computation of genomic features

#### distance to TAD border

It defines the distance of each gene's transcriptional start site (TSS) to the border of the closest topologically associated domain (TAD), where the TAD annotation from Hi-C data on mESC is taken from Dixon et al. (Dixon et al. 2012).

#### distance to Xist

It defines the linear distance of each gene's TSS to the TSS of the Xist gene (Gencode Version M9 gene annotation on mm10 was lifted over to mm9).

#### distance to LADs

It defines the distance of each gene's TSS to the closest Lamina Associated Domain (LAD) boundary. The genomic annotation of LADs in mESCs was taken from Peric-Hupkes et al. (Peric-Hupkes et al. 2010).

#### overlap with LADs

It indicates whether a region of 1000 bp around each gene's TSS overlaps or not with the annotation of a LAD. LADs annotation in mESC was taken from Peric-Hupkes et al. (Peric-Hupkes et al. 2010). This feature is dichotomic: a value of "1" indicates an overlap of the gene's 1000 bp region with a LAD, "0" indicates no overlap.

#### gene density

It is defined by the number of annotated genes within the 200 kb region around each gene TSS. Gene annotation is taken from gencode version M9 on mm10 and lifted over to mm9.

#### overlap with Xist early sites

Engreitz et al. defined the genomic coordinates of few early site (between 100 kb and 1 MB in size) on the X chromosome, which have been identified as regions coated by Xist at an early stage of XCI, i.e. sites where Xist transfers itself from its transcription locus in order to initiate spreading across the X chromosome (Engreitz et al. 2013). We compute the overlap of each X-linked gene with these early sites and define a dichotomic feature where a value of "1" indicates an overlap between the gene and an early site, while "0" indicates no overlap.

#### HiC 3D interactions

number of 3D interactions or strength of 3D interactions (sum(HiC interactions strength) / number of interactions) defined by HiC data for each gene's promoter. Interactions are subdivided into all interactions (*number interactions (HiC) all* and *mean interaction strength (HiC) all*), interactions with other promoters only (*number interactions (HiC) promoter* and *mean interaction strength (HiC) promoter)* or with the Xist locus (*mean interaction strength (HiC) xist*). HiC data was taken from Schoenfelder et al. (Schoenfelder et al. 2015).

#### HiCap 3D interactions

HiCap is a technique which combines Hi-C with sequence capture of promoter regions, so it identified promoter-anchored 3D chromatin interactions at high-resolution. We compute three features from HiCap data on mESCs (Sahlén et al. 2015): *number interactions (HiCap) all*, which corresponds to the total number of interactions of each gene's promoter with other elements, such as other promoters or enhancer regions, averaged over two replicates; *number interactions (HiCap) promoter*, which corresponds to the number of interactions of each gene's promoter with other promoters only and *number interactions (HiCap) enhancers*, which corresponds to the number of interactions of each gene's promoter with enhancer elements only.

#### overlap with CpG islands

It indicates whether a region of 1000 bp around each gene's TSS overlaps or not with an annotated CpG island. The CpG island annotation was taken from UCSC on mm9. This feature is dichotomic: a value of "1" indicates an overlap of the gene's 1000 bp region with a CpG island, "0" indicates no overlap.

#### CpG content

It defines the normalized CpG content within the 1000bp region around each gene's TSS, computed as the ratio of observed over expected CG dinucleotides (Marsico et al. 2013): $\frac{\#GpGs\ /\ L}{((\#G + \#C)\ /\ 2L)^2}$ where $L$ is the length of the considered region.

### Supplemental Text 2: Clustering details

#### Determination of the optimal number of clusters

We developed a scoring system for determining the optimal number of clusters *k* for the forest-guided clustering procedure described in the main text. We chose the optimal number of clusters by minimizing the model bias while restricting the model variance. The model bias usually measures how far off the real model with a certain *k* value is from the expected model is, while the variance is related to model complexity: complex models have high variance and poor generalization capability.

We define the model bias by the $mixture\_index_k$ which penalizes values of *k* yielding a clustering with a high degree of mixture (i.e. clusters containing genes from both silencing classes). For the definition of the $mixture\_index_k$ we introduce a "mixture" measure for each cluster *i* which is defined as:

$$mixture\_index_i \; = 4 * (\frac{x_{i0}}{n_i} * \frac{x_{i1}}{n_i})$$

where $n_i$ is the number of genes in cluster *i* and $x_{ij}$, with either $j = 0$ *or* $j = 1$, is the number of genes from cluster *i* belonging to silencing class *j*. The maximum value of the mixture for each cluster *i* is 0.25 in case of a very mixed cluster where 50% of genes belong to one class and 50% to the other class. We multiply $mixture\_index_k$ by a scaling factor of 4 to obtain a number between 0 and 1. A small adjustment to this formula is needed in case of class imbalance. The smaller class needs to be scaled to the size of the larger class in a way that both classes have comparable influence on the index value. Hence, the number of genes belonging to the smaller class $x_{sj}$ in cluster *i* are scaled by:

$$scaled\ x_{sj} = x_{sj} + \frac{x_{sj}}{n_{small}} * (n_{large} - n_{small})$$

where $n_{small}$ is the total number of genes belonging to the smaller class and $n_{large}$ is the total number of genes belonging to the larger class.

The $mixture\_index_k$ for a given number of clusters $k$ represents the average degree of mixture per cluster across all $k$ clusters:

$$mixture\_index_k = \frac{\sum_{i=1}^{k} mixture\_index_i}{k}$$

The smaller the value of the $mixture\_index_k$ the better the separation of both class into separate clusters. On the other hand we restrict the model variance to discarding too complex models and thereby avoid overfitting. Therefore, we analyse the "stability" of the forest-guided clustering for each value of $k$. We assess the stability of each cluster in the clustering by resampling the data 300 times via bootstrapping and then computing the Jaccard Similarity for each cluster. The Jaccard Similarity for each cluster is defined as:

$$JS(A|B) = \frac{|A \cap B|}{|A \cup B|}$$

where A is the set of genes in the original cluster and B is the set of genes in the same cluster after bootstrapping the data. The analysis is performed with the function clusterboot of the R package fpc. Jaccard similarities values which are smaller or equal to 0.5 are an indication of a "dissolved cluster", while values higher than 0.6 are usually indicative of stable patterns in the data (Hennig 2008). We define a clustering to be stable if each cluster in the partition has a Jaccard Similarity (JS) > 0.6. Only stable clusterings, i.e. clustering with low variability, are considered as clustering candidate for selecting an optimal value of $k$ based on the minimal bias. Hence, the optimal number of clusters $k$ is the one yielding minimum $mixture\_index_k$ while having a stable clustering.

In addition, we only consider partitions with $k > 2$ in order to unravel the combinatorial rules from the Random Forest Model.

#### Optimal number of clusters for both models

Considering the scoring system described above, the optimal number of clusters for the XCI/escape model is $k = 4$, with stable clusters (Jaccard similarities > 0.6) for each cluster (see **Supplementary**

**Figure 7**). The minimal value of $mixture\_index_k$ for the silencing dynamics model is $k = 3$, which also has a stable clustering (Jaccard Similarities > 0.7) for all clusters (see **Supplementary Figure 7**).

### Supplemental Figures

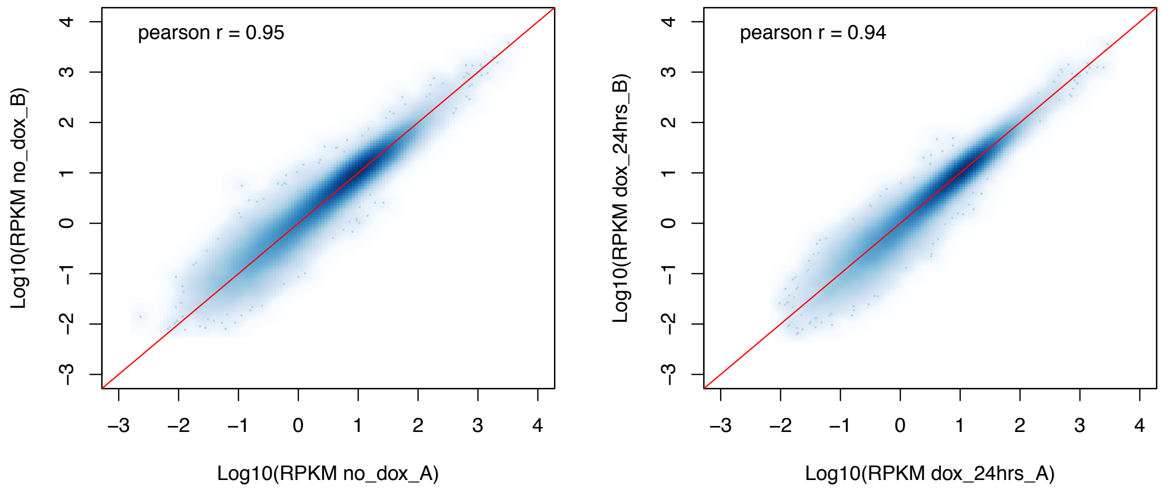


**Supplemental Figure 1: Correlation of PRO-Seq replicates.** Scatterplots of the log10 RPKM of all autosomal genes of (**a**) no doxycycline sample A and B and (**b**) dox 24 hrs sample A and B.

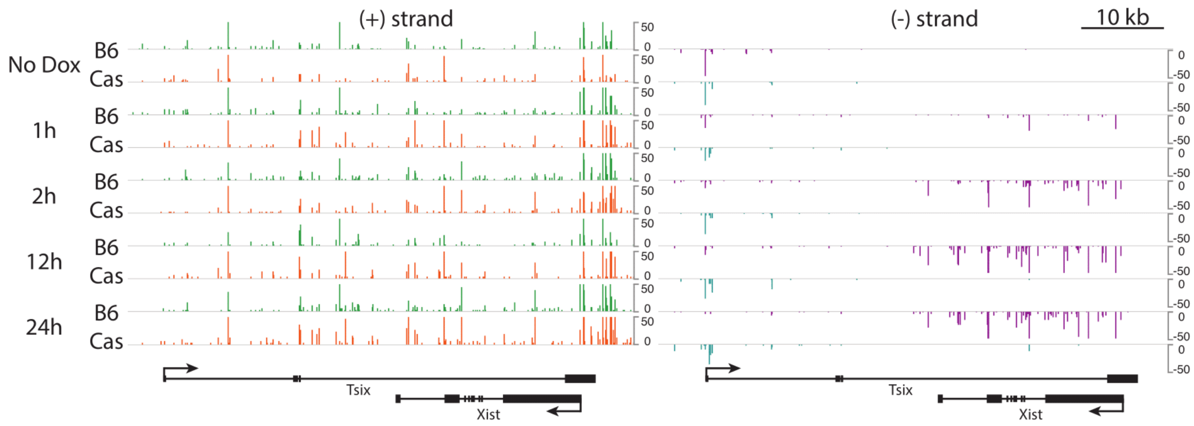


**Supplemental Figure 2: Xist Tsix region allelic track.** Tracks of normalized allelic stranded PRO-seq reads at the Xist and Tsix gene in time. On the left the (+)-strand, with the Black6 track in green and the Castaneus track in orange. On the right the (-)-strand, with the Black6 track in pink and the Castaneus track in cyan.

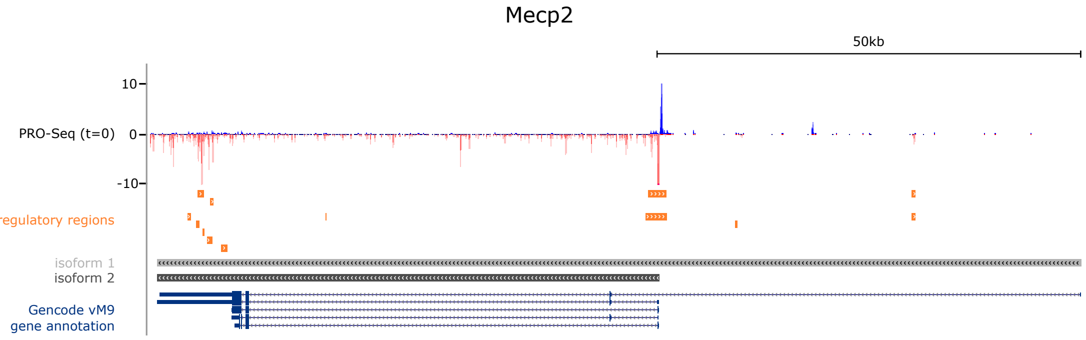


**Supplemental Figure 3: Example of assignment of an X-linked gene to its active TSS.** Gencode vM9 gene annotation and annotated regulatory regions from the PRO-seq data (no doxycycline), identified with the dREG tool (Danko et al. 2015) are used to assign each gene its corresponding active promoter/TSS. As example, the Mecp2 gene on the (-) strand of chromosome X is shown. Its assigned active TSS is the one corresponding to isoform 2, as it overlaps a regulatory region defined by a bi-directional peak in the PRO-Seq track.

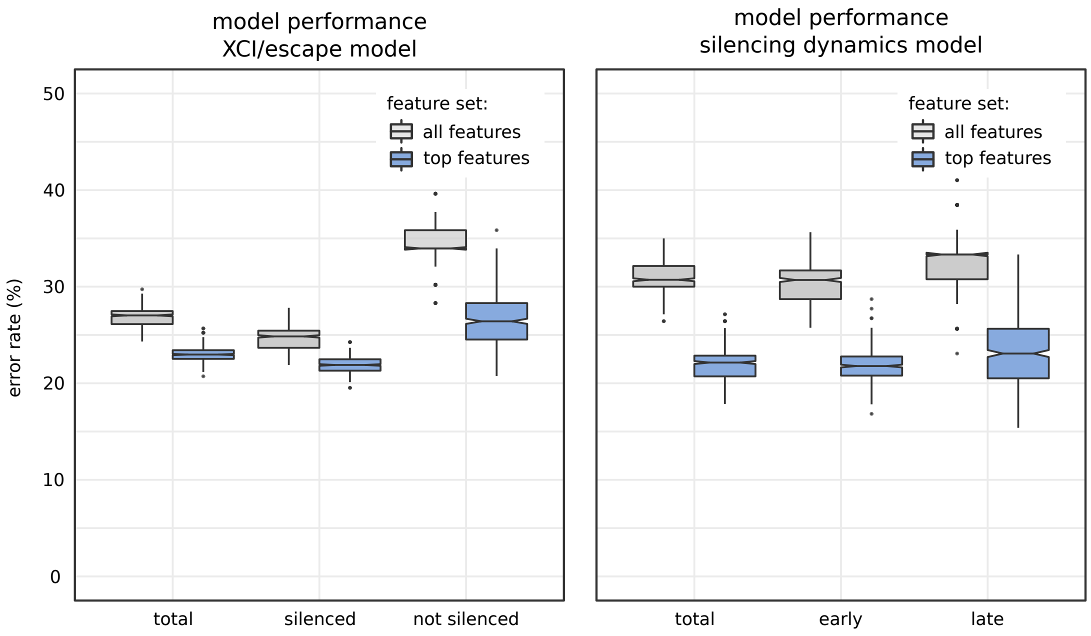


**Supplemental Figure 4: Model performance.** Random Forest (RF) model performance measured from the Out-of-Bag error rate (as defined in the Methods) for the XCI/escape model (left panel) and the silencing dynamics model (right panel). Each box in the plot represents the distribution of error rates over 500 trained RF models. Error rates are reported for both classes combined ('total') and for the prediction of each individual class (silenced and not silenced class for the first model, early and late silenced class for the second model). In addition, error rates are reported for models trained on the complete set of features (74 epigenetic and genomic features, namely 'all features'), as well as for the best models trained only on the 'top features' according to Random Forest variable importance analysis (**Supplementary Figure 5** and **Supplementary Figure 6**).

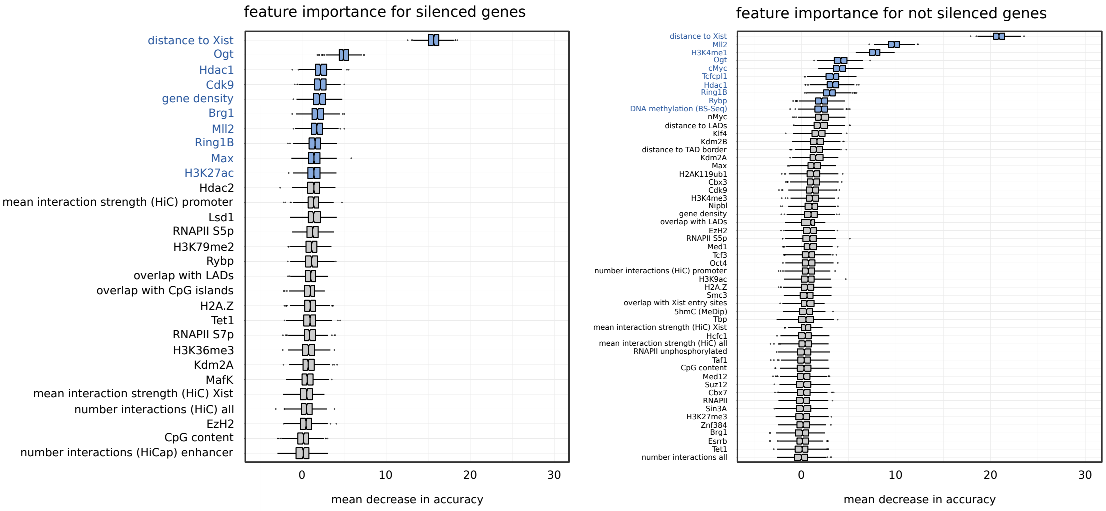


**Supplemental Figure 5: Feature importance for the XCI/Escape model.** The importance of features for Random Forest classification is measured by the mean decrease in accuracy (MDA), which is defined as the average decrease in model accuracy from permuting the values in each feature. The feature with the highest MDA (e.g. distance to Xist) is the most important feature for classification. Each box in the plot corresponds to a model feature and represents the distribution of that feature's MDA over 500 Random Forest models. For simplicity, only features with MDA higher than 0 are shown. Features marked in blue are the top selected features (10 in the XCI/escape model) which are used for final classification.

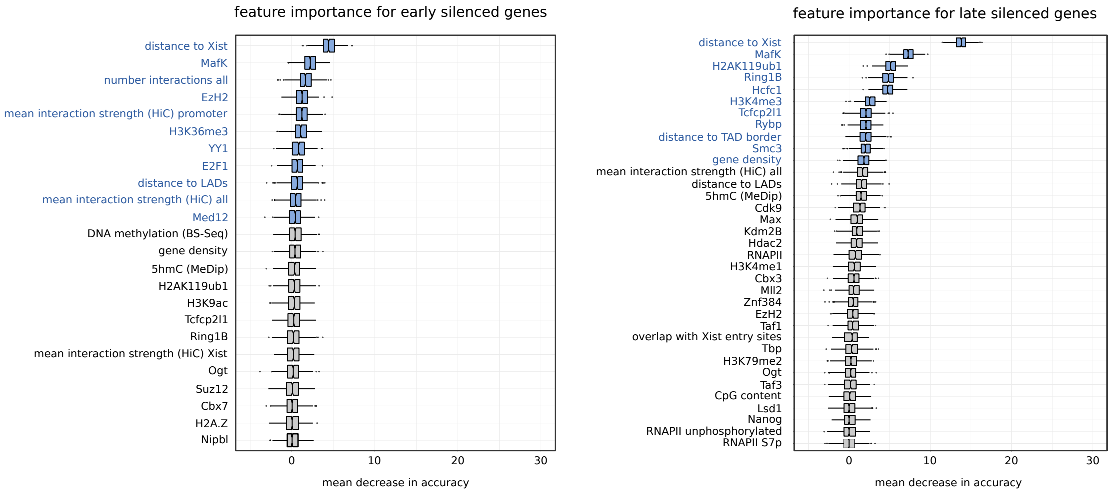


**Supplemental Figure 6: Feature importance for the silencing dynamics model.** The importance of features for Random Forest classification is measured by the mean decrease in accuracy (MDA), which is defined as the average decrease in model accuracy from permuting the values in each feature. The feature with the highest MDA (e.g. distance to Xist) is the most important feature for classification. Each box in the plot corresponds to a model feature and represents the distribution of that feature's MDA over 500 Random Forest models. For simplicity, only features with MDA higher than 0 are shown. Features marked in blue are the top selected features (11 in the silencing dynamics model) which are used for the final classification.

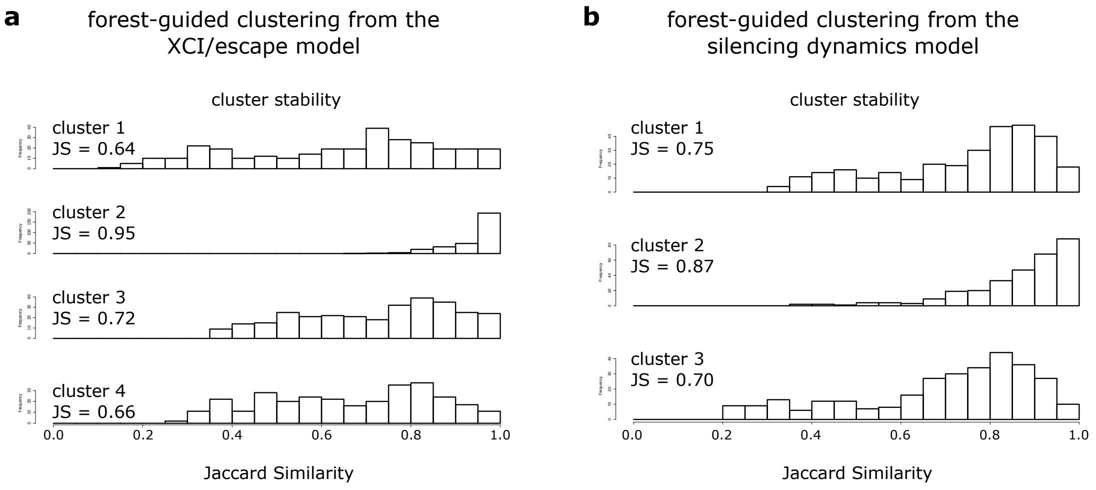


**Supplemental Figure 7: Cluster stability analysis for optimal number of *k* clusters.** The cluster stability analysis shows the distribution of Jaccard Similarity (JS) for each cluster over 300 bootstrap runs. Average JS values over 300 runs are reported for each cluster. (a) Cluster stability of XCI/Escape model for *k=4*. (b) Cluster stability of silencing dynamics model for *k=3*.

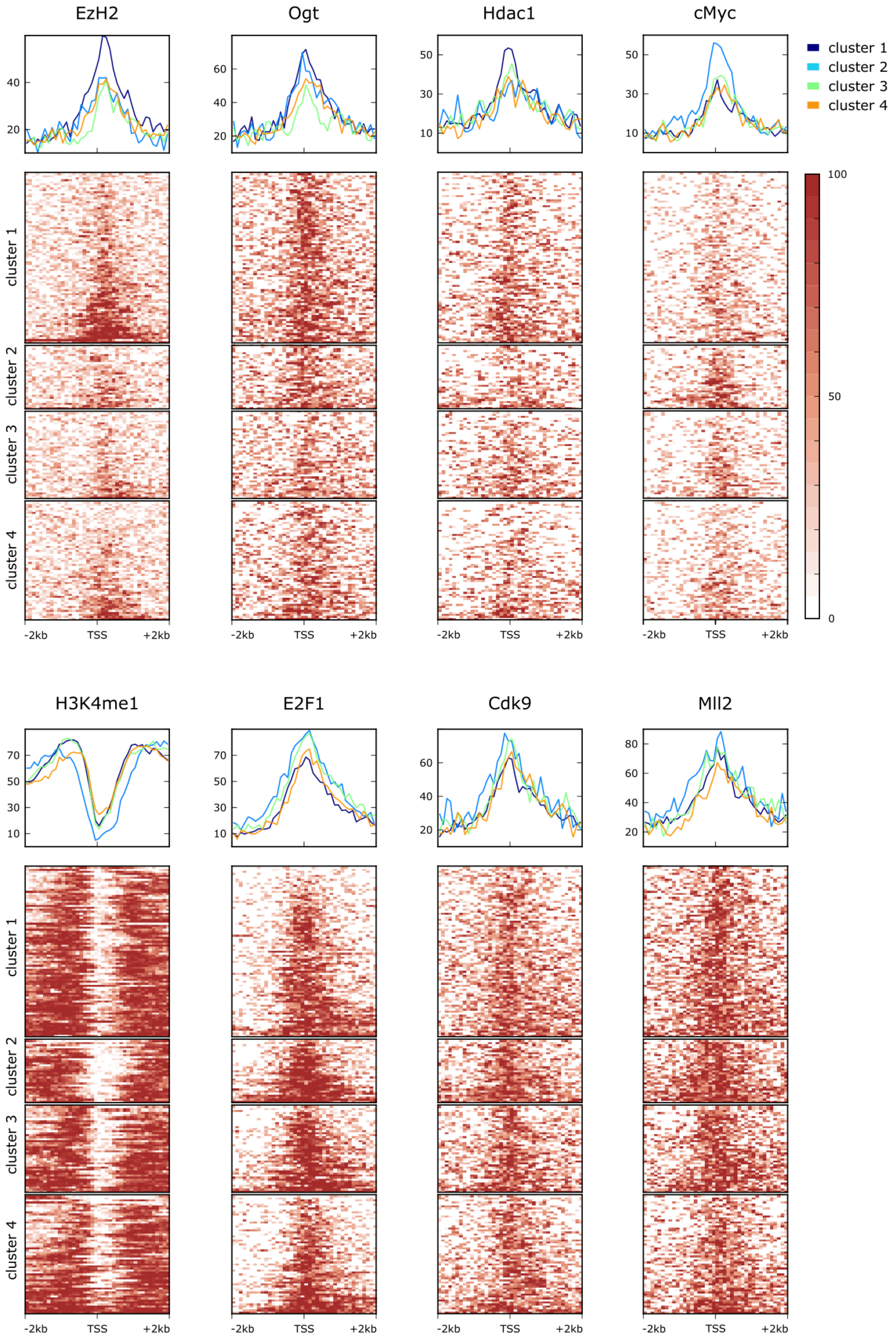


**Supplemental Figure 8: Enriched features from the XCI/escape model clustering.** The normalized signal of epigenetic marks and other factors computed in the +/- 2000 bp genomic region around 280

X-linked gene promoters is shown in the heatmaps for each of the four clusters separately. Average profile plots for the same factors are also shown above the heatmaps to highlight overall differences between clusters. Shown here are only those features which, according to the p-value of an ANOVA test, were the top 10 most significantly different among clusters in the XCI/escape model.

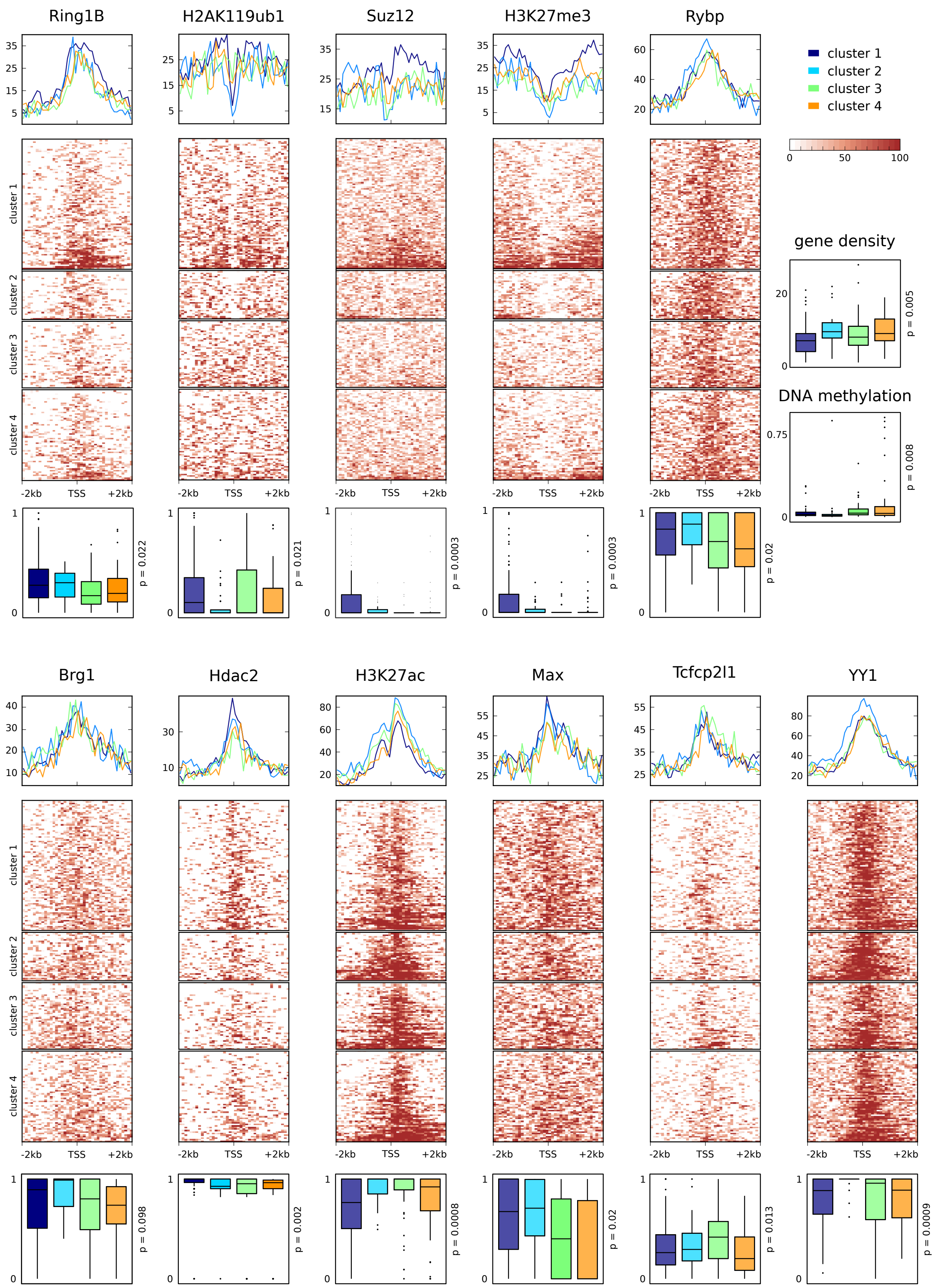


**Supplemental Figure 9: Top features from the XCI/escape Random forest.** The normalized signal of epigenetic marks and other factors, computed in the +/- 2000 bp genomic region around 280 X-linked gene promoters is shown in the heatmaps for each of the four clusters separately. Average profile plots for the same factors are also shown above the heatmaps to highlight overall differences between clusters. Shown here are epigenetic and genomic features that are among the top features in the XCI/escape Random Forest model (**Supplementary Figure 5**) but are not among the top 10 significant ones from the clustering. Boxplots of feature distributions in each cluster are shown below the heatmaps to give an idea of their variation among clusters.

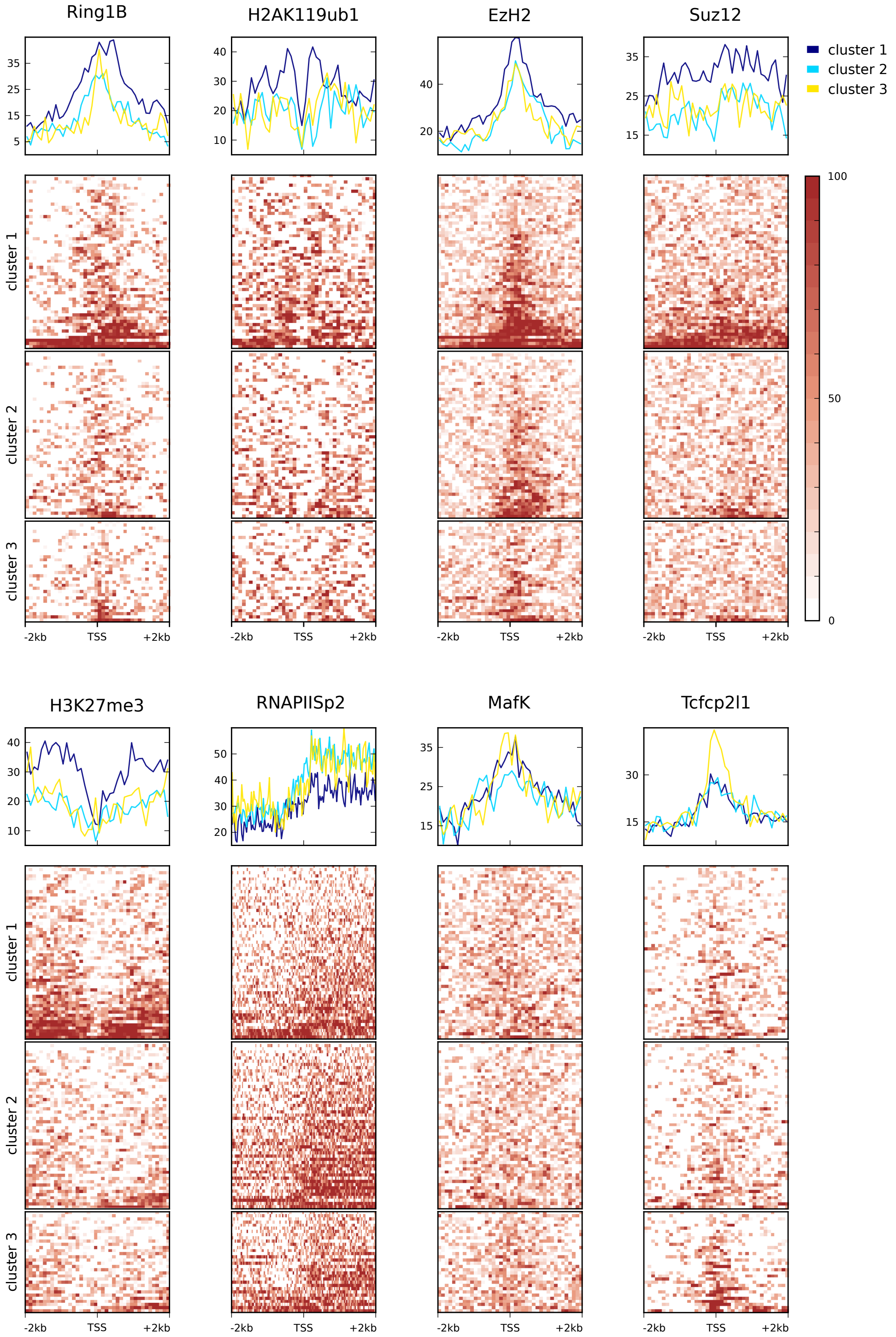


**Supplemental Figure 10: Enriched features from the silencing dynamics model clustering.** The normalized signal of epigenetic marks and other factors, computed in the +/- 2000 bp genomic region around 280 X-linked gene promoters is shown in the heatmaps for each of the four clusters separately. Average profile plots for the same factors are also shown above the heatmaps to highlight overall differences between clusters. Shown here are only those features which, according to the p-value of an ANOVA test, were the top 10 most significantly different among clusters in the silencing dynamics model.

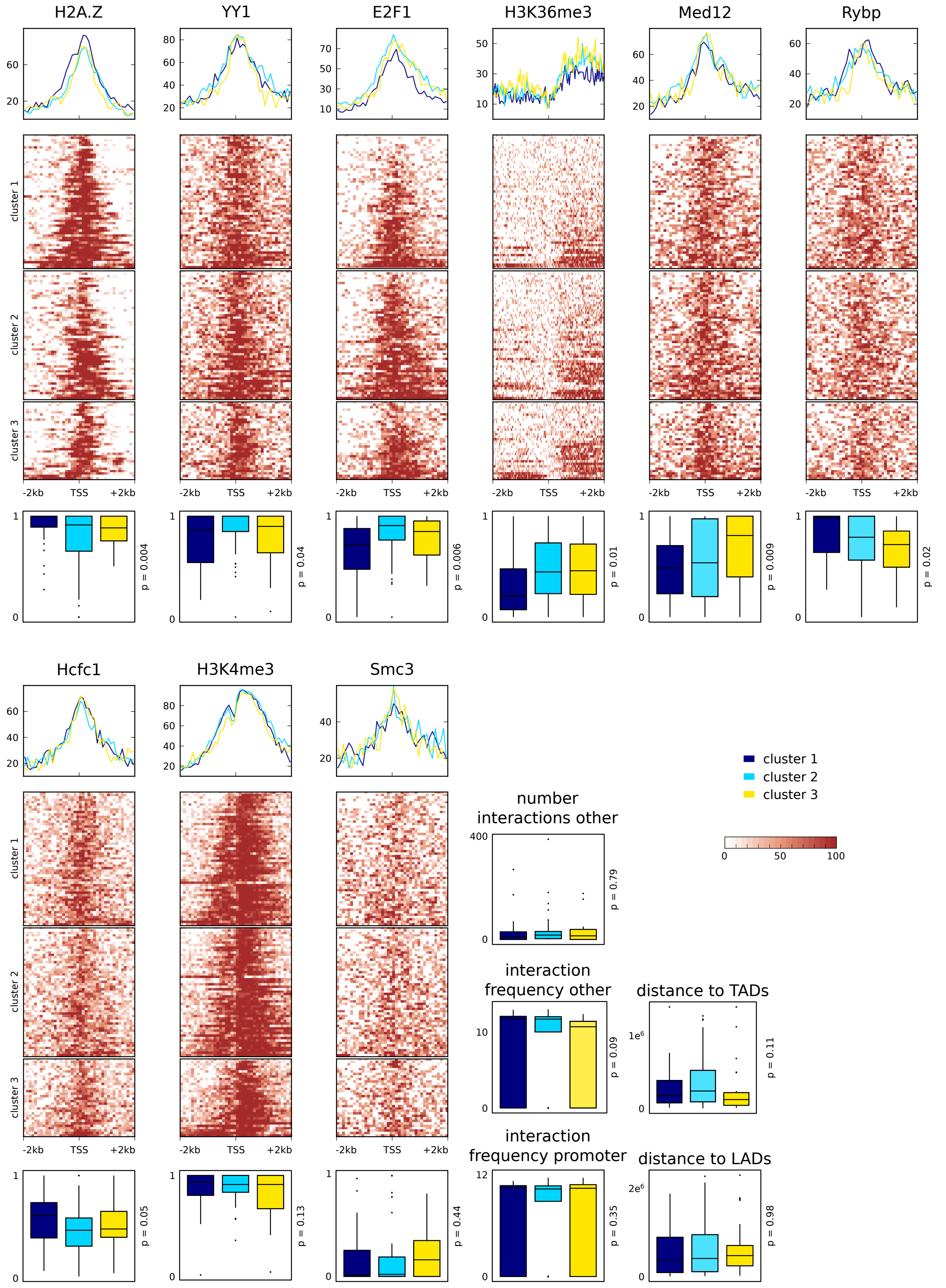


**Supplemental Figure 11: Top features from the silencing dynamics Random forest.** The normalized signal of epigenetic marks and other factors, computed in the +/- 2000 bp genomic region around 280 X-linked gene promoters is shown in the heatmaps for each of the four clusters separately. Average profile plots for the same factors are also shown above the heatmaps to highlight overall differences between clusters. Shown here are epigenetic and genomic features that are among the top features in the silencing dynamics Random Forest model (**Supplementary Figure 6**) but are not among the top 10 significant ones from the clustering. Boxplots of feature distributions in each cluster are shown below the heatmaps to give an idea of their variation among clusters.

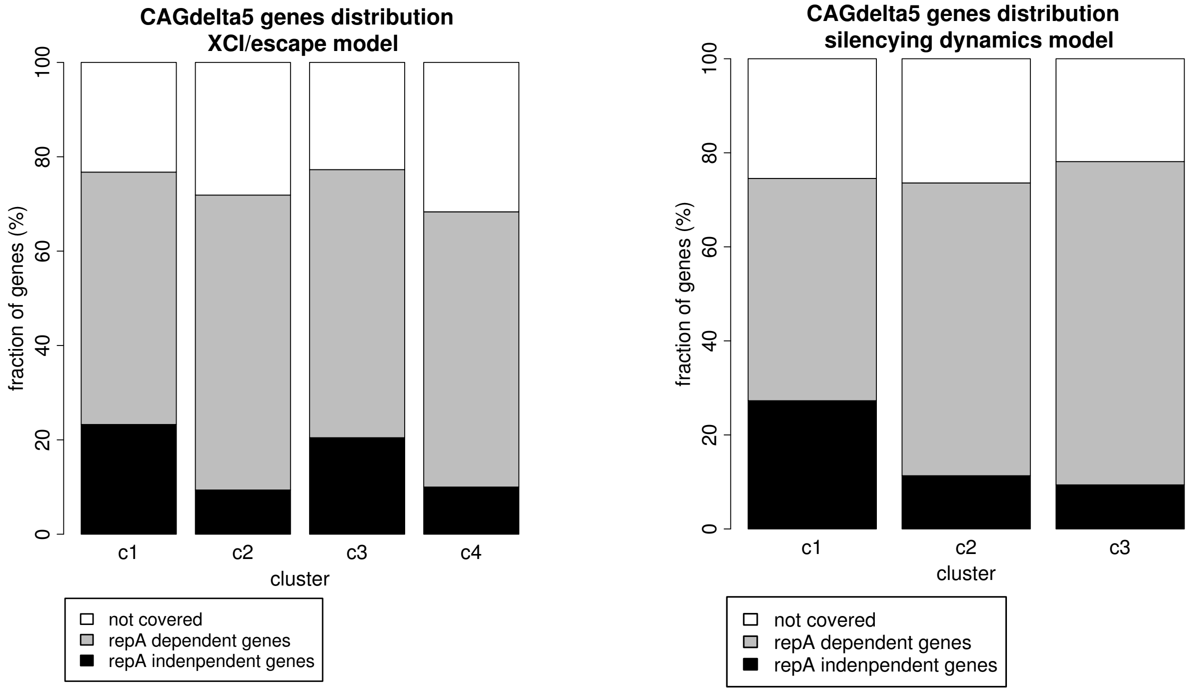


**Supplemental Figure 12: Comparison of clustering results from both models with classification of genes in repeat A-dependent and repeat A-independent from Sakata et al. (Sakata et al. 2017).** The proportion of genes shown to undergo silencing in mouse trophoblasts independent or dependent of Xist repeat A element is shown for each cluster for the XCI/escape model (left panel) and the silencing dynamics model (right panel). In detail, 'repA dependent genes' refers to those genes from (Sakata et al. 2017) which showed abrogated silencing in Xist-repeat A-mutant-cells, 'repA independent genes' refers to those genes which could still undergo silencing in the same cells,and 'not covered' refers to those genes in our dataset which were not covered in the Sakata et al. study (Sakata et al. 2017).

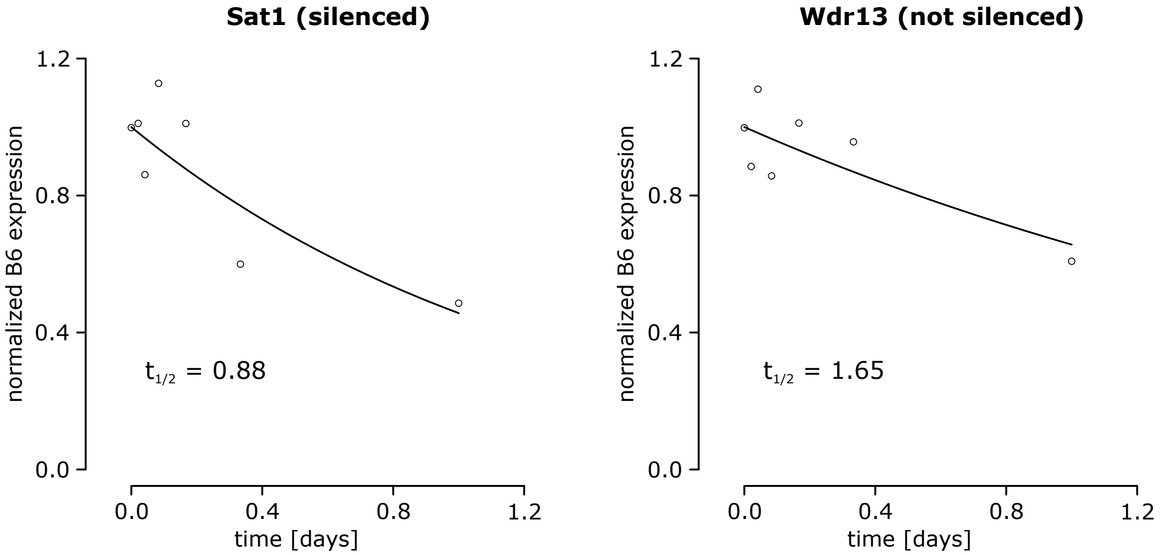


**Supplemental Figure 13: Experimental validation of Sat1 and Wdr13 with Pyrosequencing.** Experimental validation of one gene predicted as silenced (Sat1) and another gene predicted as not silenced (Wdr13) with Pyrosequencing. The experiment was conducted over 24 hours with 7 time points. We normalized the B6 expression and computed the half-time (as defined in the Methods). The half-time of Sat1 (0.88) lies within the range of silenced genes (< 0.9) and the half-time of Wdr13 (1.65) lies within the range of not silenced genes (> 1.6), as correctly predicted by the XCI/escape model.

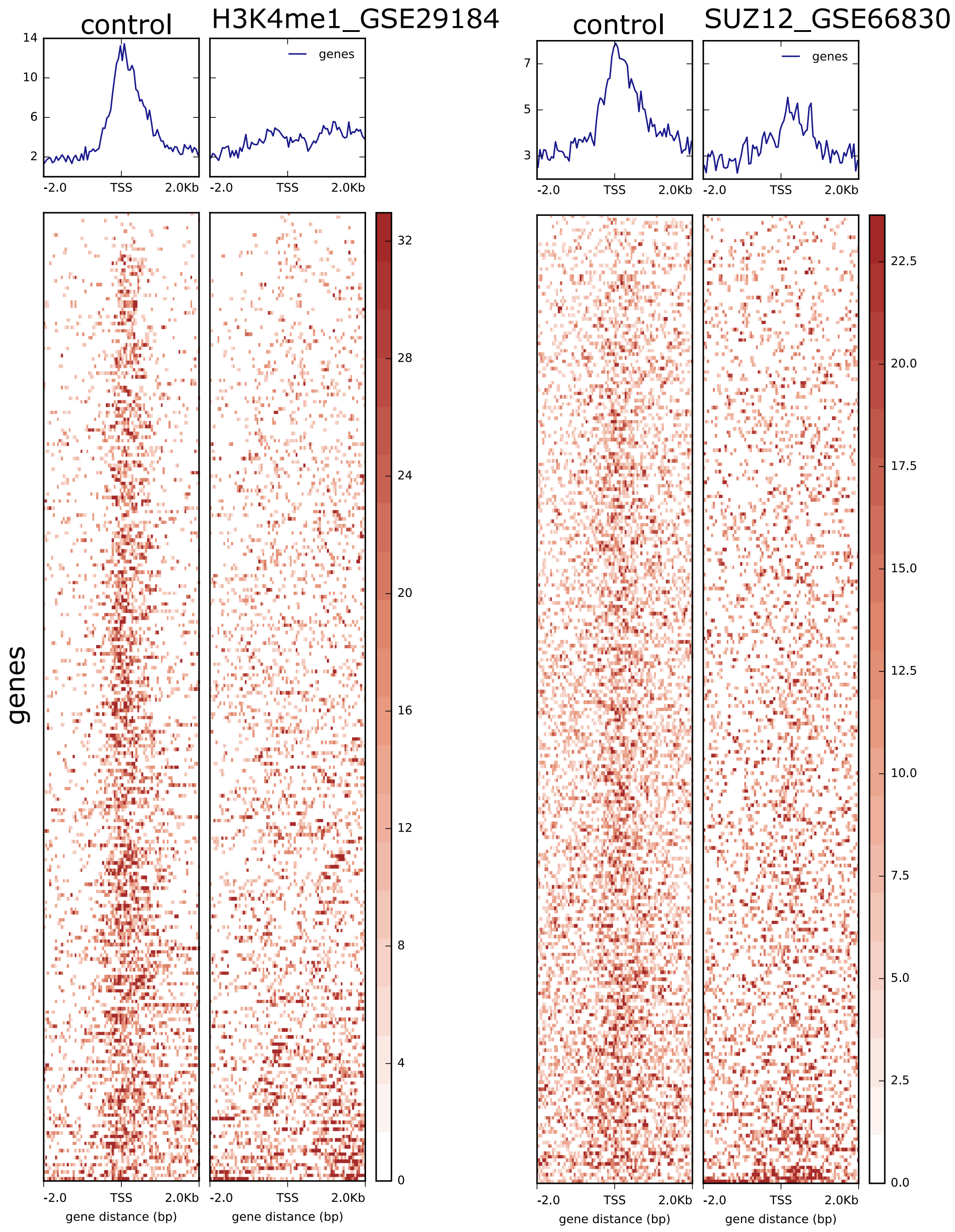


**Supplemental Figure 14: ChIP-Seq library filtering with deepTools heatmap.** DeepTools heatmaps are visualized for two ChiP-Seq experiments, H3K4me1 (GEO: GSE29184, left panel) and SUZ12 (GEO: GSE66830, right panel) and their respective input controls. Shown is the ChIP-seq signal at the +/- 2000bp region around the TSS of each gene (280 X chromosomal genes with computed half-times).

Both experiments show a higher enrichment for the control input signal in this region than the experiment itself (where very little signal is present). This evidence makes the quality of both data sets doubtful and therefore those libraries, as well as other data sets showing similar characteristics, were excluded from further analysis to avoid biases in the modelling process.

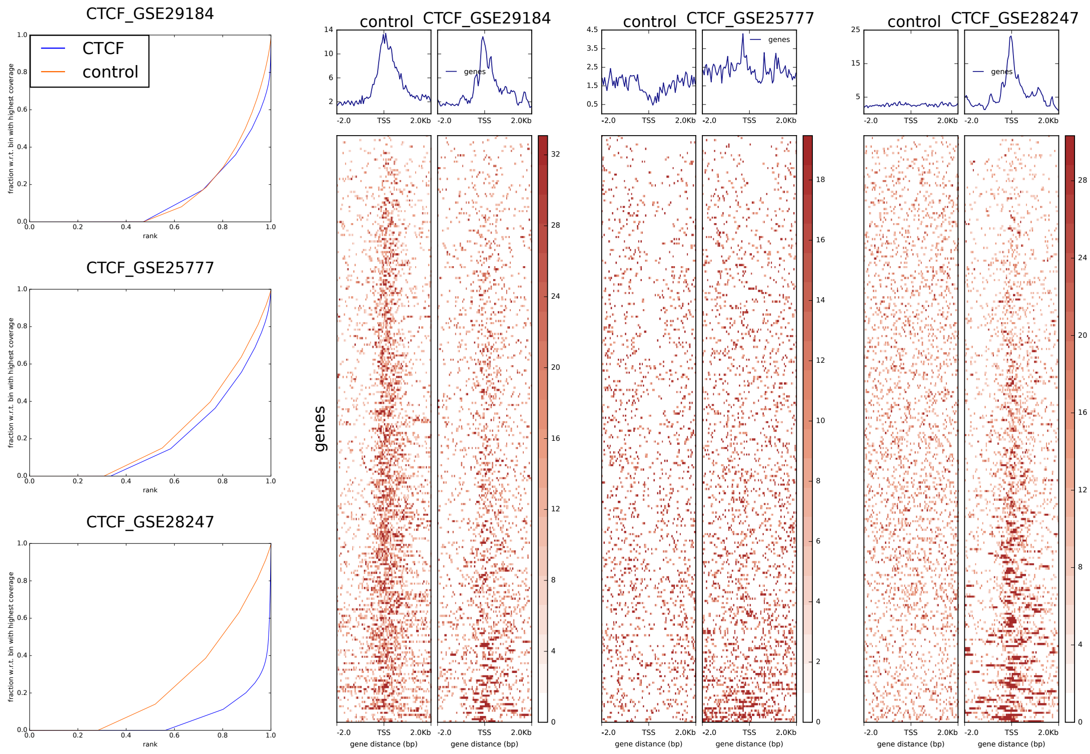


**Supplemental Figure 15: Example on how to select the best ChIP-seq data set for a given factor.** When more than one data set was available in GEO for a given factor or epigenetic feature only one is selected for further analysis based on deepTools heatmaps (on the right) and fingerprint plots (on the left). An example is shown for CTCF, where the dataset with GEO GSE28247 is selected out of three libraries as 1) the fingerprint plot shows that the cumulative distribution of the reads from the input experiment (orange line) is closer to the diagonal, indicative of a uniform read distribution, compared to the other two libraries; 2) the fingerprint plot shows that the read distribution of the ChIP experiment (blue line) has a steep rise toward the end of the plot, it clearly distinguishes itself from the control read distribution (orange line), and it is therefore indicative of a nonuniform, but rather peaked, read distribution at CTCF binding sites, compared to the other two libraries; 3) the heatmap clearly shows signal enrichment for CTCF at the -/+ 2000 bp region around gene TSSs compared to input, indicative of a good signal to noise ratio, which is not the case for the GSE29184 and the GSE25777 libraries.

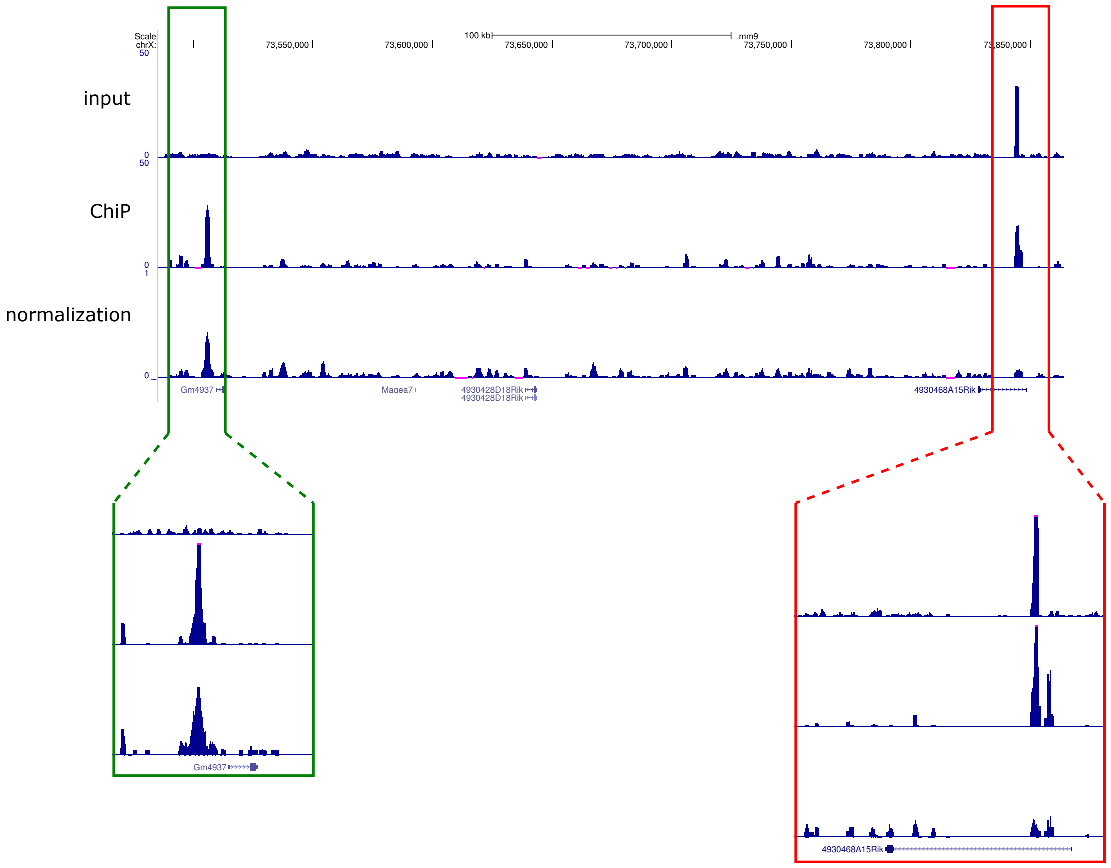


**Supplemental Figure 16: Example of ChIP-Seq signal normalization with normR.** Two genomic loci on chromosome X are shown. The green box highlights a region with no or little uniform signal in the input control but a sharp peak in the ChIP library. The normalized track correctly shows that the signal corresponding to the sharp peak is still maintained after normalization. In contrast, the red box highlights a region with a peak signal in both the input control and the ChIP library. The normalized track correctly shows that the peak in this region is rescaled after normalization to the input signal.

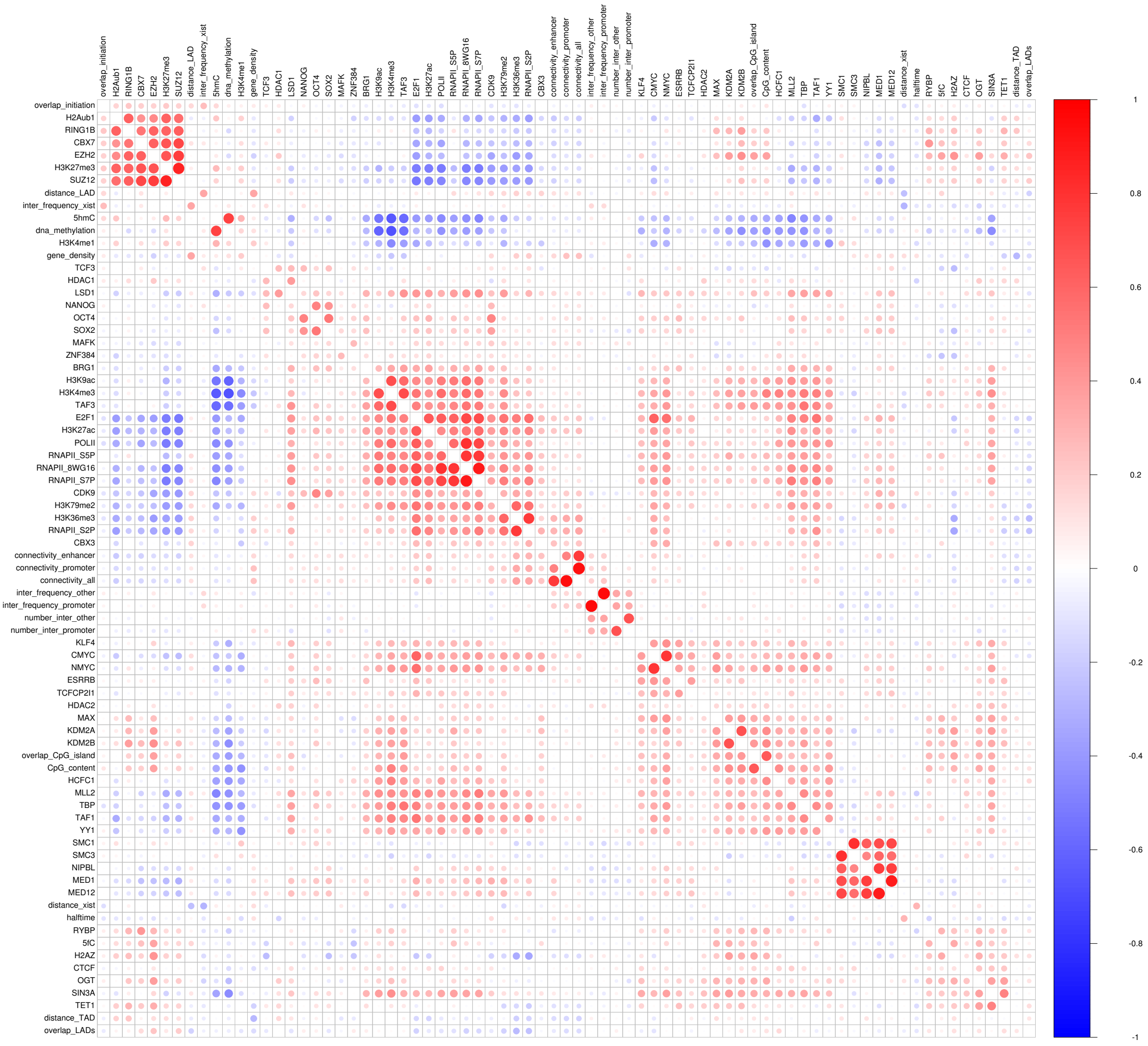


**Supplemental Figure 17: Feature correlation matrix**. It shows the Pearson correlation for every pair of features used in the model and it is computed based on all 280 genes with estimated half-times from the PRO-seq data. Red indicates high positive correlation and blue a high negative correlation. One can observe blocks of correlate features. For example, the active marks (PolII, H3K4me3, H3K27ac and others) are highly correlated amongst each other while repressive features, such as PRC1 and PRC2 components and H3K27me3 form another positively correlated block but are negatively correlated with many active mark features.

### Supplemental Tables

**Supplemental Table 1: List of Random Forest predictions for genes without half-times.** Sheet 1 contains a list of candidate genes for experimental validation. Those genes were confidently predicted as silenced (class 0) or not silenced (class 1) and were selected based on prediction probability, expression at time point 0 and number of SNPs falling into their exons (see Material and Methods for further information). Sheet 2 contains the whole predictions list for all genes without computed half-times. Predicted classes marked in red are not reliable. The additional columns list the value of each feature in the XCI/escape model for each gene.

**Supplemental Table 2: List of genes with computed half-times.** This table contains 280 genes for which we could measure half-times. The genes are listed by gene name with corresponding genomic position (annotation of active TSS, see Material and Methods). In addition to the half-time column, there are columns listing the basal skewing (basal skewing towards one allele at t=0), sqrtRSS (fitting error of the exponential decay function), RPKM value of each gene at t=0, if a gene is a known escapee shown by other studies (source given in brackets with legend at the top of the gene list) and the class predicted by our Random Forest model for each gene after training averaged over 500 RF models.

**Supplemental Table 3: Metadata of Random forest features.** Sheet 1 lists all ChIP-Seq, BS-Seq and genomic data used for the Random Forest model with its source and a description of the feature. For ChIP-Seq data the enrichment region (upstream and downstream of the TSS) is given. Sheet 2 contains information about the filtering of ChIP-Seq data and the reason why certain libraries were discarded.

**Supplemental Table 4: Filtering steps in the computation of half-times.**

| **filtering step** | **# of genes after filtering** |
|---|---|
| no filter | 484 |
| minimum read coverage per timestep > 10 | 341 |
| basal skewing between 0.2 and 0.8 | 330 |

| | |
|---|---|
| sqrtRSS < 1.5 | 296 |
| regulatory region within defined region around TSS | 280 |

**Supplemental Table 5: Filtering steps in the ChIP library pre-processing.**

| filtering step | # of ChIP libraries after filtering |
|---|---|
| no filtering | 133 |
| remove ChIP libraries with < 3 Mio. reads | 120 |
| manuel filtering with heatmap plots | 79 |
| selecting the best ChIP library for each feature | 57 |

**Supplemental Table 6: Ranges of half-times for choosing class thresholds.**

| class | half-times ranges |
|---|---|
| silenced genes | $t_{1/2} < [0.9,\ldots,1.4]$ |
| not silenced genes | $t_{1/2} > [1.4,\ldots,2]$ |
| early silenced genes | $t_{1/2} < [0.5,\ldots,0.7]$ |
| late silenced genes | $[0.7,\ldots,1] < t_{1/2} < [1,\ldots,1.4]$ |
